# An Unusual Follower Peptide is Required for Biosynthesis of the Antibiotic Lasso Peptide Triculamin

**DOI:** 10.64898/2026.07.03.736388

**Authors:** Tiziana Svenningsen, Aske Merrild, Asger Buchbjerg Petersen, Aliana Nosolini Dos Reis, Anne Mette Pold, Hannah Lange, Thomas Tørring

**Affiliations:** Department of Biological & Chemical Engineering, Aarhus University, Gustav Wieds Vej 10C, 8000, Aarhus, Denmark

**Author notes:** Co-first authors (Tiziana Svenningsen, Aske Merrild).

**Keywords:** heterologous expression, lasso peptides, non-canonical biosynthesis, RiPPs, triculamin

## Abstract

Triculamin is a potent antibiotic lasso peptide first isolated in 1967. Previous studies have demonstrated that its biosynthesis follows a non-canonical logic unlike any other lasso peptide. In this study, we investigate the role of the unusual follower peptide and demonstrate that it is essential for efficient biosynthesis. Using structural prediction and targeted mutations of key conserved residues, we hypothesize that the interactions between the follower peptide and the macrocyclase create an enzyme-substrate complex that ensures delivery of the core peptide to the enzyme active site. Moreover, we demonstrate that analogs of the lasso peptide can be produced by modifying the core peptide, highlighting the substrate promiscuity of the lasso macrocyclase and identifying lysine-3 in the lasso peptide ring as the site of acetylation. Lastly, we achieve successful heterologous expression in *Burkholderia sp*. FERM 3421, which proves to be a superior heterologous host.

## Introduction

Lasso peptides are a structurally and biosynthetically distinct class of ribosomally synthesized and post-translationally modified peptides (RiPPs), characterized by threaded macrolactam structure and a highly conserved biosynthetic path (1). To date most studies have been performed in so-called “canonical” systems. In canonical lasso peptides, biosynthesis proceeds from a precursor peptide comprising an N-terminal leader sequence and a core peptide, which is converted into the mature product through the coordinated action of a RiPP recognition element (RRE; B1), a leader-peptidase (B2), and an isopeptide bond-forming macrocyclase (C) (Fig. 1A) (2–4). For canonical lasso peptides, the leader peptide is well-known to function as a recognition element for the biosynthetic machinery.(5) It helps recruit and position the precursor for processing, helps leader cleavage by the B2 protein, and subsequent macrolactam formation by the C protein (6). Conserved features in the leader, such as the penultimate threonine in many lasso peptide precursors, have also been shown to be important for recognition and processing (7,8).

**Figure 1.**
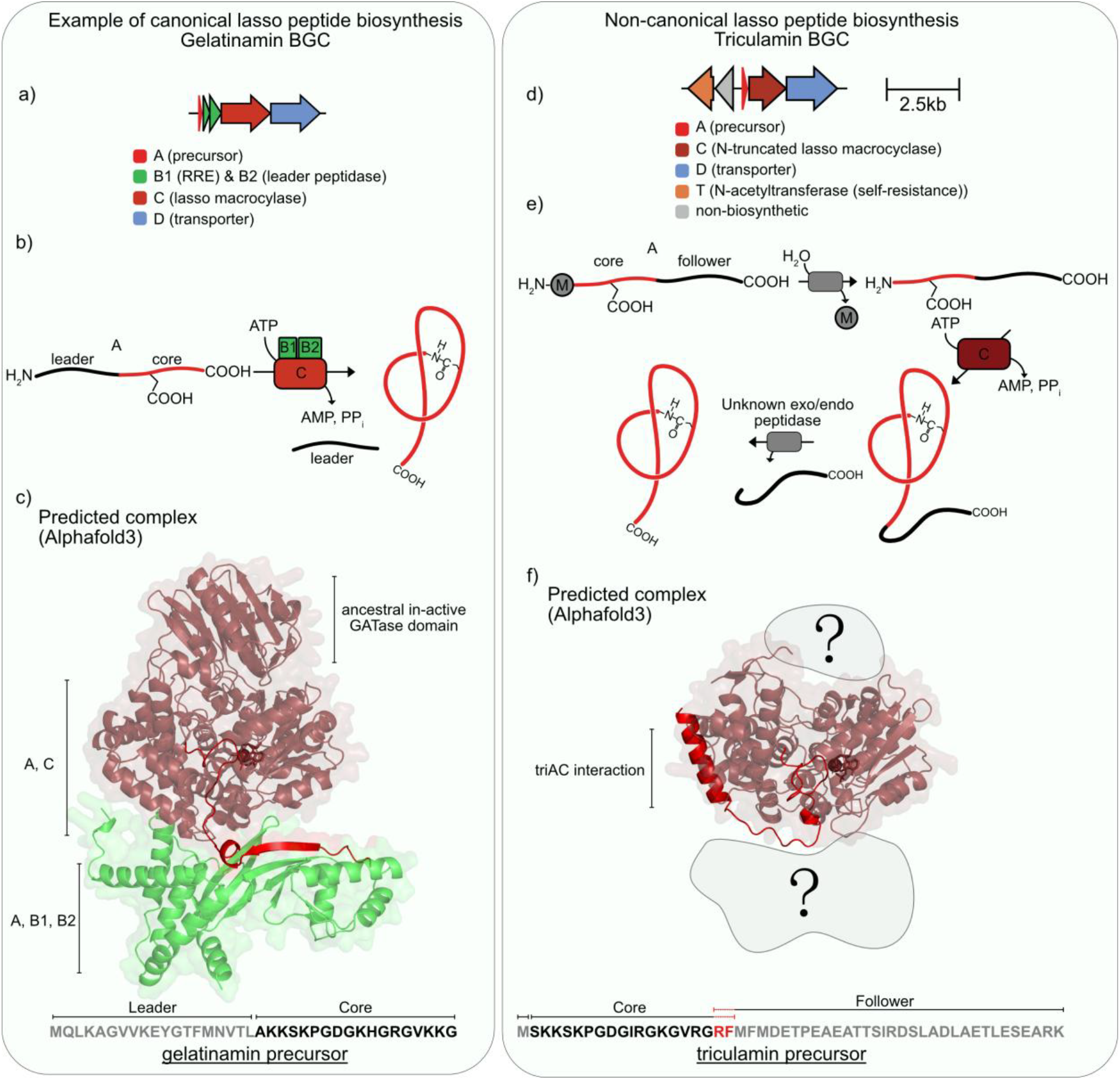
Schematic of canonical and non-canonical lasso peptide biosynthesis. a) The lasso peptide gelatinamin is from a canonical BGC containing the precursor peptide, B1/B2 processing enzymes, C macrocyclase, and D transporter. b) Canonical lasso peptide biosynthesis, where the leader peptide is recognized and cleaved by the B proteins, followed by ATP-dependent macrocyclization of the core peptide by the C enzyme. c) Alphafold3 (23) predicted complex of the gelatinamin biosynthetic enzymes (GelAB1B2C), where the precursor is shown light red. d) Triculamin represents a non-canonical lasso peptide BGC, lacking the canonical B proteins and instead encoding the precursor peptide, TriC macrocyclase, TriD transporter, and TriT N-acetyltransferase. e) Hypothesized triculamin maturation pathway. The precursor contains an N-terminal core and a C-terminal follower sequence, which is removed during biosynthesis. TriC catalyzes macrocyclization, while TriT acetylates the mature peptide. f) Alphafold3 (23) predicted complex of triAC suggesting an interaction between the follower and macrocyclase itself.

In previous work, we described, what appeared to be a “non-canonical” lasso peptide biosynthesis of the lasso peptide triculamin.(9,10) Triculamin, and the identical alboverticillin, initially discovered in the 1950s-1960s as an antimycobacterial peptide antibiotic produced by *Streptomyces triculaminicus* and *S. griseocarneous*,(11,12) follows a biosynthetically atypical biosynthesis that in contrast to all other characterized lasso peptides, is produced from an unusually compact biosynthetic gene cluster (BGC) that encodes only a precursor peptide, a transporter, a truncated macrocyclase (Fig. 1f), and an *N*-acetyltransferase. Notably, this macrocyclase lacks the conserved ∼100–250 amino acid catalytically inactive N-terminal domain, which is responsible for glutamine amidotransferase activity in the ancestral asparagine synthetase (Fig. 1c,f). Even more strikingly, the triculamin precursor lacks a canonical N-terminal leader peptide and instead contains a C-terminal follower sequence (9). This feature had not been observed in lasso peptides prior to our previous studies (9,10), and rare among other bacterial RiPP classes, with few comparable example reported, exemplified by bottromycin (13). These major differences in biosynthetic pathways conceal these non-canonical BGCs from algorithms which bioinformatically predict lasso peptide BGCs as the missing genes are essential BGC annotation (14,15).

Motivated by these deviations from the canonical paradigm, we subsequently investigated the prevalence and biosynthetic diversity of triculamin-like peptides. This analysis revealed that these peptides are more widespread than previously appreciated and occur in both canonical and non-canonical lasso peptide BGCs. Heterologous expression studies demonstrated that closely related core peptides can be matured via distinct biosynthetic routes: triculamin exemplifies a non-canonical pathway, whereas palmamin and gelatinamin are produced through canonical systems (10,16).

At present, little is known about how the triculamin system achieves precursor recognition, proteolytic processing, and macrocyclization. The conservation of follower sequences across diverse non-canonical triculamin-like BGCs suggests that these elements may play an active role in biosynthesis. Understanding the non-canonical biosynthetic strategies will provide nuances on the mechanistic limits of lasso peptide formation, but it also begs the question as to why all canonical lasso peptide biosynthetic pathways employ a precursor-RRE-peptidase complex only to remove the leader as the first step of the biosynthesis. Early work on Microcin J25 reported that the macrocyclase, MccjC, cannot form the mature lasso peptide from the core sequence alone (17). Curiously, conflicting data has been reported on whether minute amounts of a synthetic core peptide can be processed when incubated with both MccjB and MccjC.(18,19) Similar results have been observed for fuscanodin and particularly pseudomycoidin which shows efficient biosynthesis in absence of the leader, RRE and peptidase (20–22).

In this study, we use heterologous expression of the triculamin BGC in *Streptomyces albidoflavus* to investigate whether the follower sequence function analogously to leader peptides in canonical lasso peptide biosynthesis. We further assess whether specific residues within the follower sequence are especially important for *in vivo* biosynthesis. Moreover, we perform heterologous expression in *Burkholderia sp*. FERM-3421 and *E. coli* BL21(DE3) to determine the compatibility of the triculamin BGC with heterologous hosts outside the Actinomycetota.

## Results and discussion

### The C-terminal follower sequence is required for efficient triculamin biosynthesis

Through exploration of triculamin and its biosynthesis, we have been interested in reconstituting the macrocyclase *in vitro*. In this regard, we attempted *E. coli* heterologous expression of triC, purification and reconstitution, unsuccessfully. Thus, we directed our attention to performing mutations in the triculamin BGC in the functioning heterologous expression system of *S. albidoflavus*.

Work on canonical lasso peptide systems has shown that precursor peptides often require only minimal sequence elements for productive maturation. For example, in the microcin J25 (MccJ25) system, only a short region of the leader peptide is necessary for processing, with specific residues acting as docking motifs for the biosynthetic enzymes rather than contributing to bioactivity (17). These findings suggest that precursor peptides primarily encode recognition elements for maturation machinery.

In our previous work, we bioinformatically determined that all precursor peptides found in non-canonical BGCs contain a long follower peptide that is absent from the final natural products (9,10). Using AlphaFold3(23) to predict the structural interaction with the macrocyclase, two observations are worth noting: 1) in all cases examined, the follower is predicted to fold into an alpha helix, 2) while the core peptide is predicted to enter the active site of the macrocyclase, the follower helix shows high-confidence interaction with the macrocyclase surface. Based on these observations, we hypothesized that genetically removing the helix or parts of it would disrupt the precursor-macrocyclase interaction and be detrimental to biosynthesis of triculamin in the *S. albidoflavus* host. We therefore designed a series of follower peptide truncations to test this hypothesis. In total, 11 variants were constructed in which the follower sequence was progressively shortened in increments of three amino acids, yielding a series of constructs lacking between 3 and 33 residues of the C-terminal region (Fig. 2). The cloning workflow for generating all mutations can be seen in Fig S3. Each variant was expressed under identical conditions, and triculamin production was quantified by LC-mass spectrometry. Quantification was estimated based on an MS standard curve from purified triculamin A (Fig. S1 and S2).

**Figure 2.**
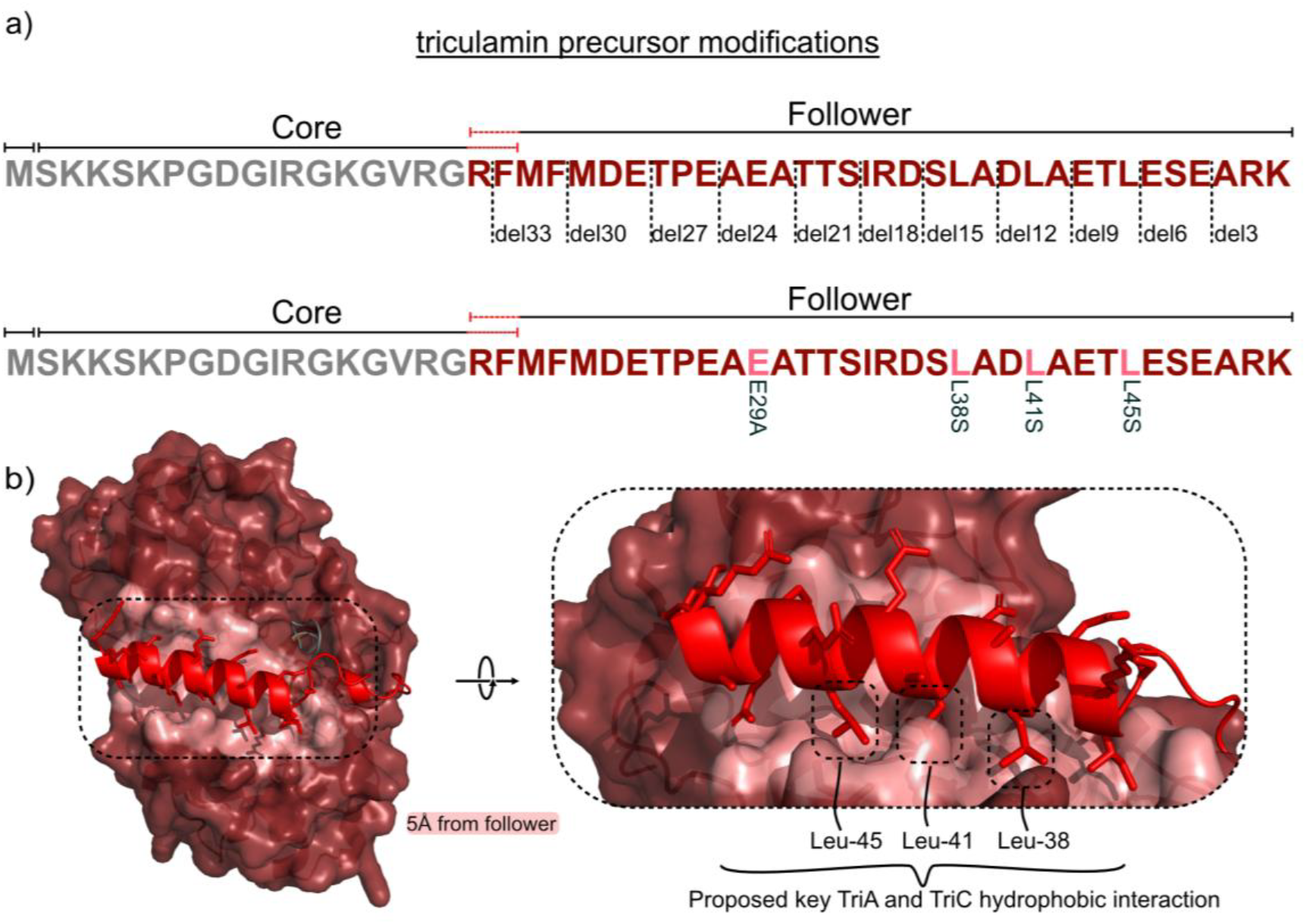
Triculamin precursor modifications and proposed follower–protein interactions. a) Overview of the triculamin precursor variants used to probe the follower region. The core peptide is shown in grey and the C-terminal follower in red. Deletion variants and selected point mutations are indicated relative to the precursor sequence. b) AlphaFold3 model shows the follower peptide interacting in with a hydrophobic pocket of triC. Residues Leu-38, Leu-41, and Leu-45 are positioned toward a hydrophobic surface patch, suggesting that these residues contribute to key follower-mediated recognition. Salmon color indicates surfaces within 5 Å of the follower.

Deletion of the terminal three amino acids resulted in a slight increase in triculamin production relative to the full-length precursor, although this difference was not statistically significant. In contrast, removal of six or more amino acids led to a pronounced reduction in product formation. For all truncations beyond this point, no triculamin could be accurately detected by LC-MS.

These results demonstrate that nearly the entire C-terminal follower sequence is required for efficient triculamin biosynthesis. The detrimental effects observed by even modest truncations suggest that the follower sequence plays a critical role. Based on the AlphaFold3 prediction, this could indicate that the follower helix ensures stronger affinity for the macrocyclase and thereby increases catalytic efficiency, but it could also merely be that the extended tail increases stability against proteolytic degradation and thereby the intracellular half-life (24).

### Triculamin biosynthesis is abolished by mutating conserved amino acids of the follower sequence

Inspection of the triculamin precursor revealed three conserved leucine residues within the C-terminal follower sequence that are positioned to interact with the macrocyclase (Fig. 2b). These leucine residues are also conserved across many previously identified non-canonical triculamin-like precursor peptides, suggesting that they may be functionally important for follower-mediated recognition (10).

Structural analysis of the predicted complex suggested that these residues project toward a hydrophobic pocket in the macrocyclase, raising the possibility that hydrophobic interactions mediate precursor recruitment to the macrocyclase. This is analogous to the canonical leader recognition, in which hydrophobic regions of the precursor’s leader segment drive interactions with the RRE (7,8). To test this hypothesis, we individually substituted each of the three leucine residues with serine, thereby disrupting hydrophobic side-chain interactions. In addition, double, and triple mutants were constructed in which two out of three and all three leucine’s were simultaneously replaced by serine. Each variant was expressed under identical conditions, and triculamin production was assessed by LC-MS.

Mutation of any single leucine residue resulted in a marked reduction in triculamin production relative to the wild-type precursor (Table 1). This effect was further exacerbated in the triple mutant, for which product formation was reduced drastically.

**Table 1.** Effect of follower-region deletions and point mutations on total triculamin production in fermentation broth. triA refers to the wild-type precursor peptide and served as the reference for statistical comparisons of all truncation and mutation variants. Total triculamin is the sum of triculamin A, A-Ac, B, B-Ac, C, and C-Ac. Concentrations were estimated from MS signal using a standard curve generated with purified triculamin A and are reported as mean ± SD. Statistical significance was assessed using raw two-sided Welch t-tests relative to *triA(WT)*. Trace indicates signal below the calibrated 0.1 μg/mL reporting limit; ns, not significant; *, p < 0.05 ^a^.

| Follower Variants | Total Triculamin<br>Broth concentration (μg/mL, mean ± SD) |
| --- | --- |
| *triA*(WT) | 1.7 ± 0.5 |
| *del3* | 3.9 ± 1.7 ns |
| *del6* | trace (<0.1) * |
| *del9 – del33* | Not detected |
| *E29A* | Not detected |
| *L38S* | 1.7 ± 0.4 ns |
| *L41S* | 0.6 ± 0.2 * |
| *L45S* | 0.1 ± 0.0 * |
| *L41S/L45S* | Not detected |
| *L38S/L41S/L45S* | 0.1 ± 0.1 * |

These results indicate that the conserved leucine residues within the follower sequence are critical for efficient triculamin biosynthesis. The strong sensitivity of the leucine-to-serine substitution supports a model in which hydrophobic interactions between the follower sequence and the macrocyclase contribute directly to precursor recognition. In addition to the leucine we included a mutation, E29A. The rationale for this mutation was a high degree of conservation across the triculamin family. Changing the aspartate to alanine completely abolished production of triculamin. This highlights that conserved amino acids of the follower sequence may directly affect triculamin biosynthesis.

### TriC accepts variants in the core sequence, which reveals lysine-3 as the site of acetylation

Previous studies of triculamin had shown that one of three lysine residues in the core peptide (position 2, 3 or 5) is acetylated for self-resistance (10). However, MS/MS analysis did not allow unambiguous assignment of the acetylation site, and NMR-based structural characterization was not possible due to limited quantities of an acetylated variant. To further analyze both the functional role of these residues and the substrate tolerance of the macrocyclase, we introduced a series of mutations within the core peptide. Variants were constructed targeting both the macrolactam ring and the tail region, including S1A and V15A, as well as substitutions at positions 2 and 3 (K2A, K2T, K3A, and K3R). All variants were expressed under identical conditions, and product formation and modifications were analyzed by LC-MS.

The mutation in the tail (V15A) and within the ring (S1A) were well tolerated, with triculamin production observed at levels comparable to the wild-type precursor (Fig. 3b). In contrast, substitutions at positions 2 and 3 had more pronounced effects. The K2A and K3R variants exhibited substantially reduced production. K2A produced only acetylated triculamin, whereas K3R produced only non-acetylated. Interestingly, the K3A variant yielded high levels of triculamin with no acetylated products (Fig. 3b). That all mutations were accepted indicates that the triculamin macrocyclase exhibits a degree of tolerance to mutations in both the ring and tail region of the core peptide. Additionally, this opens the possibility of generating analogs with improved characteristics. The observation that no acetylation is found in the K3A, and K3R variants suggests that K3 may be the main site of acetylation. The mutant K2T shows both acetylated and non-acetylated triculamin, supporting this observation. Detailed MS/MS analysis of triculamin B-Acetylated, compared to the K2A B-variant acetylated, unambiguously identifies K3 as the acetylation site (Fig. 3c-d). Interestingly, lysine is also present at position 3 in two other triculamin-like lasso peptides that are acetylated; however, both gelatinamin and lariocidin are acetylated specifically on their tail K16 (10,25). Multiple lysine residues are likely critical for bioactivity, and thus self-resistance can be achieved by acetylation of different lysine’s.

**Figure 3.**
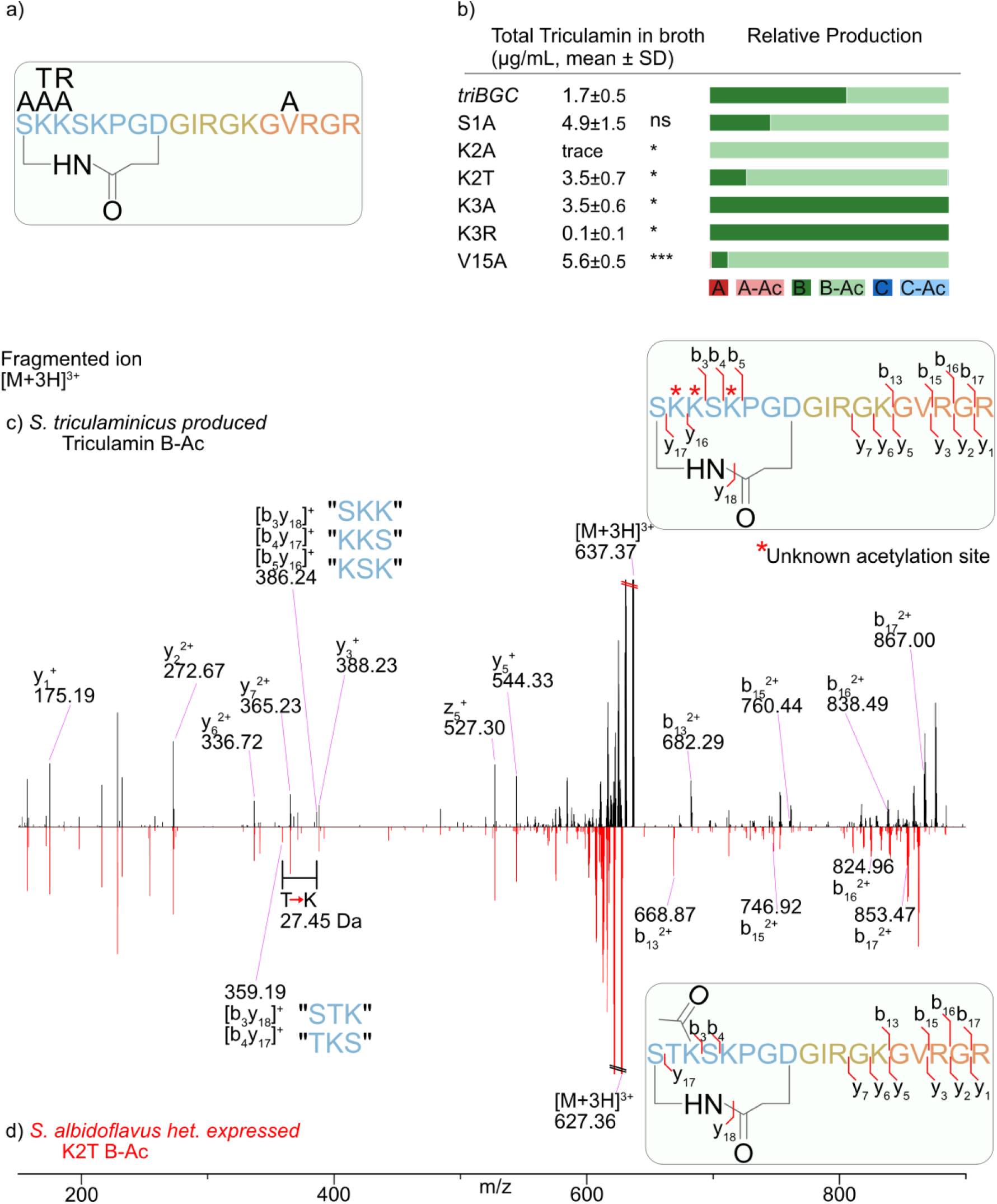
Core variants reveal K3 acetylation in triculamin a) Schematic overview of triculamin B shown as a non-lasso structure for simplicity together with designed point mutations (in black). b) Total triculamin production in fermentation broth for triA core mutants. Total triculamin was calculated as the summed abundance of triculamin A, B, and C and their acetylated forms, and broth concentrations were estimated from MS signal intensities using a triculamin A standard curve. Values are shown as mean ± SD. Colored bars indicate the relative distribution of the detected triculamin variants within each mutant. Statistical significance was assessed using two-sided Welch’s t-tests against the total triculamin production in *S. albidoflavus*; * p < 0.05, ** p < 0.01, *** p < 0.001. c) MS/MS mirror plot of triculamin B-Ac from *S. triculaminicus* plotted against the mutant triculamin K2T B-Ac heterologously expressed in *S. albidoflavus*. The same y-fragments are observed, whereas b-fragments correspond to the lysine to threonine mutation. Double fragmentation of the lasso ring distinguishes K3 as the acetylation site.

### Triculamin biosynthesis proceeds in *Burkholderia*, a non-*Streptomyces* heterologous expression host

The unique and absent elements of the triculamin-BGC, in comparison with those of established lasso peptide biosynthesis, are not well understood. We wondered whether our previously unsuccessful *in vitro* reconstitution was due to a missing helper protein present in *Streptomyces* but not in the *E. coli* from which we attempted to purify the cyclase. To test this, we performed heterologous expression of the triBGC in *Burkholderia sp*. FERM-3421 Δfr9 (26,27), including the palmamin ABC-transporter (palD) (Fig. 4). Interestingly, this expression was successful and primarily produced triculamin B and C as well as acetylated variants. The same expression vector was used in *E. coli* BL21, yet it resulted in no detectable production. These results demonstrate that triculamin biosynthesis can proceed in non-*Streptomyces* hosts and that if any helper protein is required, it is shared among *Streptomyces* and *Burkholderia*.

**Figure 4.**
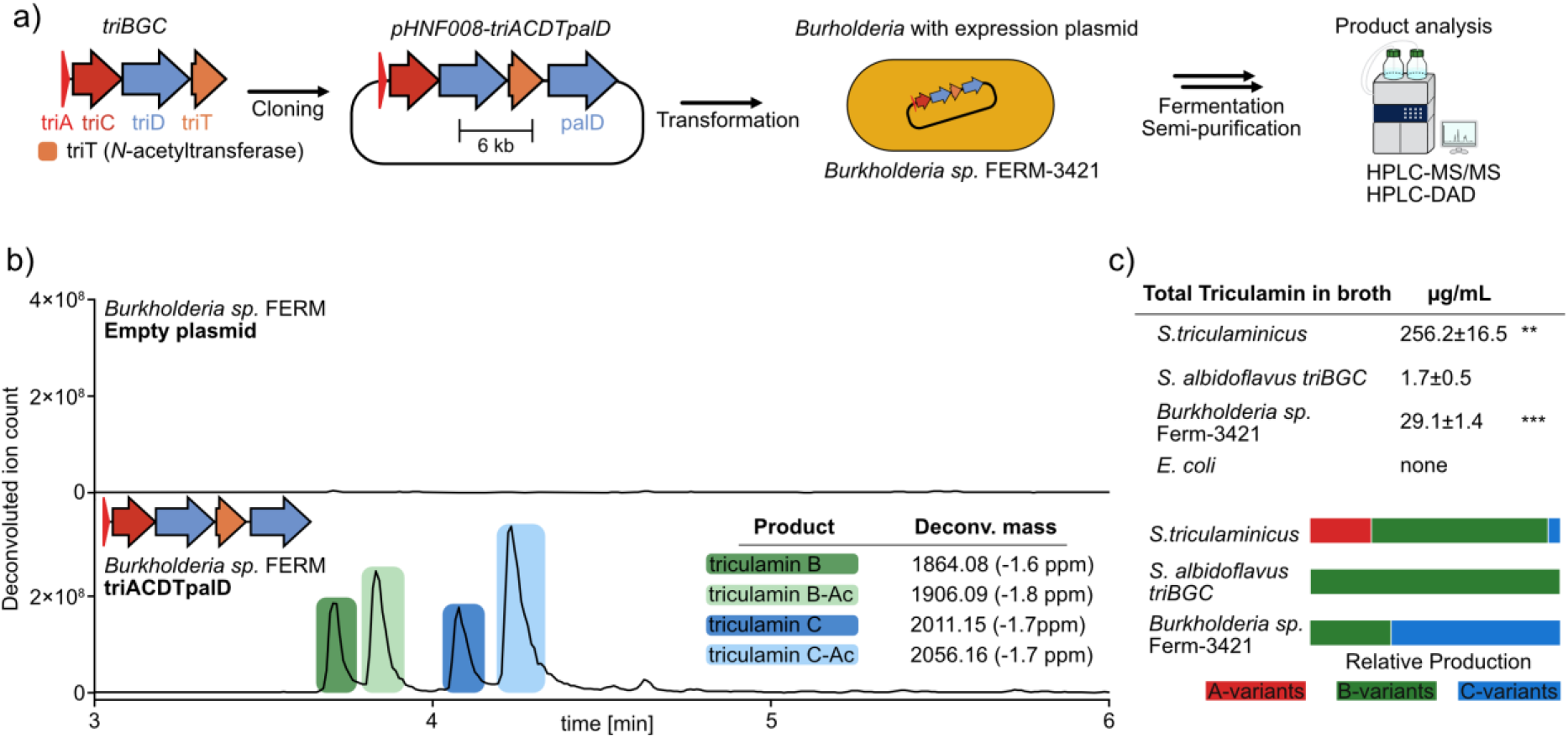
Heterologous production of triculamin variants. a) Schematic overview of the triculamin biosynthetic genes cloned into the expression plasmid pHNF008-palD to create pHNF008-triACDTpalD and introduced into *Burkholderia sp*. FERM-3421. b) Representative LC-MS chromatograms showing the empty plasmid control and *Burkholderia sp*. FERM-3421 carrying the pHHNF008-triACDTpalD plasmid. Colored regions indicate the retention windows for triculamin A, B, B-Ac and C-Ac. c) Total triculamin production in different heterologous hosts, calculated as the combined broth concentration of triculamin A, B, C, and their acetylated derivatives. Concentrations were estimated from MS signal intensities using a triculamin A standard curve acquired on the same MS system. Values are shown as mean ± SD. Stacked bars show the relative distribution of detected triculamin variants within each strain. Statistical significance was assessed using two-sided Welch’s *t*-tests against the *S. albidoflavus* heterologous control; \*\**p* < 0.01, \*\*\**p* < 0.001.

Moreover, the changes in tail lengths (triculamin A-C) indicate a non-specific follower cleavage by the expression host. This is strongly supported by the observation of triculamin A, B, and C variants from *S. triculaminicus* (Fig S4-8), predominantly triculamin B when expressed in *S. albidoflavus*, and triculamin B and C when expressed in *Burkholderia sp*. FERM-3421 (Fig S9-12). Lastly, the obvious differences in triculamin titers, together with the genetic tractability and growth rate of *Burkholderia sp*. FERM-3421, make it the superior heterologous expression host for future studies.

## Conclusion

In this manuscript, we provide further details on the non-canonical lasso peptide triculamin to confirm the necessity of the enigmatic follower peptide. We show through structural predictions that the follower likely associates with the surface of the macrocyclase. In lieu of a functioning in vitro setup, we employ heterologous production in *S. albidoflavus* J1074 to confirm that truncations as well as point mutations in conserved leucine residues are detrimental to lasso peptide formation. Furthermore, we show that the macrocyclase accepts amino acid changes in the core sequence, confirming the expected site of acetylation by the acetyltransferase TriT and opening the possibility of generating improved variants of triculamin. Finally, we show that triculamin can also be heterologously expressed in *Burkholderia* sp. FERM 3421 but primarily generates the triculamin C variants.

### Experimental procedures

Detailed experimental procedures, including media compositions, buffer formulations, primer sequences, plasmid construction, strain cultivation, purification protocols, LC-MS/MS parameters, and statistical analysis, are provided in Supplementary information.

### Strains, plasmids, and heterologous expression

The triculamin biosynthetic gene cluster and engineered *triA* variants were expressed heterologously in *Streptomyces albidoflavus* J1074. Site-directed mutations and truncations in *triA* were introduced into pL99-ACDT by PCR-based cloning followed by Gibson assembly. Sequence-verified plasmids were transferred into *S. albidoflavus* J1074 by interspecies conjugation from *Escherichia coli* ET12567. Exconjugants were cultivated in YEME medium without sucrose, and expression was induced with ε-caprolactam. Fermentation broths were harvested after five days and stored at −20 °C until analysis.

For expression outside *Streptomyces*, the triculamin biosynthetic genes were cloned into a *Burkholderia*-compatible expression plasmid together with *palD*, yielding pHNF008-*triACDTpalD*. The plasmid was introduced into *Burkholderia* sp. FERM BP-3421 Δ*fr9*, and production was performed in 2S4G medium with L-arabinose induction. The same plasmid was also tested in *E. coli* BL21(DE3).

### Sample preparation and LC-MS analysis

Fermentation broths were clarified and purified by weak cation-exchange solid-phase extraction prior to analysis. Triculamin variants were analyzed by UHPLC-DAD and HPLC-MS/MS using reversed-phase C18 chromatography. Triculamin production was quantified from deconvoluted MS signals using an external calibration curve generated with purified triculamin A isolated from *S. triculaminicus*. Total triculamin production was calculated as the summed abundance of detected triculamin A, B, C, and corresponding acetylated variants and is reported as triculamin A-equivalent concentrations.

### MS/MS annotation and acetylation-site assignment

Triculamin variants were identified by accurate mass and MS/MS fragmentation. Theoretical masses were calculated from the expected dehydrated lasso peptide structures, with acetylated variants assigned by the corresponding mass shift of +42.01057 Da. Fragmentation spectra were compared between native and heterologously produced variants to support structural assignments and determine the site of acetylation. Some spectra in the supplementary information were annotated using Interactive Peptide Spectral Annotator tool.(28)

### Structural modelling

Predicted protein–precursor complexes were generated using AlphaFold3(23) and used to guide hypotheses concerning interactions between the C-terminal follower peptide and the triculamin macrocyclase TriC. Structural models were used qualitatively to identify candidate follower residues for mutational analysis.

### Statistical analysis

Triculamin production values are reported as mean ± standard deviation from three fermentation replicates unless otherwise stated. Statistical comparisons were performed using two-sided Welch’s *t*-tests against the corresponding wildtype triculamin BGC control expressed in *S. albidoflavus*. Significance levels are indicated in the figure legends.

## Supplementary information

Supplementary information is available as a separate document Supplementary information.docx

## Acknowledgements

We gratefully acknowledge funding from the Carlsberg Foundation (CF22-1239) in the form of a Semper Ardens Accelerate grant. The authors would like to thank Alessandra Eustáquio at University of Illinois, Chicago, for sharing *Burkholderia* sp. FERM-3421 *Δfr9* and the expression plasmid pHNF008.

## Notes

### Competing Interest Statement

The authors have declared no competing interest.

