## Supplemental information for "An Unusual Follower Peptide is Required for Biosynthesis of the Antibiotic Lasso Peptide Triculamin"

#### 1 Methods

##### 1.1 General

Synthetic DNA primers were purchased from Integrated DNA Technologies (IDT), and primer sequences are listed in Table S5. PCR amplification was performed using Phusion™ Plus DNA Polymerase or DreamTaq™ Green PCR Master Mix, both obtained from Thermo Scientific. Restriction enzymes used for cloning were FastDigest enzymes from Thermo Scientific, while BsaI-HF®v2 and T4 DNA ligase used for Golden Gate cloning were obtained from New England Biolabs. DNA assembly reactions were performed using NEBuilder® HiFi DNA Assembly Master Mix according to the manufacturer's instructions.

Plasmid purification was carried out using the GeneJET Plasmid Miniprep Kit, and genomic DNA was isolated using the GeneJET Genomic DNA Purification Kit, both from Thermo Fisher Scientific. PCR products and DNA fragments were purified using the Monarch® PCR & DNA Cleanup Kit or the Monarch® DNA Gel Extraction Kit from New England Biolabs. Agarose gel electrophoresis was performed using UltraPure™ Agarose, SYBR Safe DNA Gel Stain, 6xDNA Loading Dye, and GeneRuler 1 kb DNA Ladder.

Chemically competent *Escherichia coli* DH5α cells were used for plasmid propagation. *E. coli* ET12567 was used for conjugative transfer of plasmids into *Streptomyces albidoflavus* J1074, and *E. coli* BL21(DE3) was used for heterologous expression experiments. Competent cells were prepared in-house and transformed using standard protocols. Whole-plasmid sequencing and Sanger sequencing were performed by Eurofins Genomics.

Analytical UHPLC was performed using an Agilent UHPLC system equipped with a Luna Omega Polar C18 column, as described below. HPLC–MS and MS/MS analyses were carried out on an Orbitrap Exploris™ 120 mass spectrometer coupled to a Thermo Scientific Vanquish SII LC system using electrospray ionization. Preparative HPLC purification was performed using a preparative Luna Omega Polar C18 column. MS data were processed using FreeStyle software and visualized using OriginPro 2018.

Solid-phase extraction was performed using Strata-X-CW weak cation-exchange cartridges. Media compositions and buffer formulations used in this study are listed in Tables S3 and S4, respectively.

#### 1.2 Expression and purification of triculamin from native producer *S. triculaminicus*

*Streptomyces triculaminicus* (JCM4242) was cultivated on ISP2 agar at 28 °C until sporulation. After 8 days, spores were collected from six fully sporulated plates by resuspension in 10 mL ISP2 medium per plate. Seed cultures were established by inoculating 250 mL ISP2 medium in baffled flasks and incubating at 28 °C with shaking at 200 rpm. For triculamin production, exponentially growing seed cultures were used to inoculate 2 L of triculamin production medium (TPM) in 5 L baffled flasks at 5% (v/v). Production cultures were incubated at 28 °C and 160 rpm and grown for 8 days.

Following incubation, cultures were harvested by centrifugation to remove cells. The resulting supernatant was adjusted to 50 mM HEPES (pH 8), clarified by repeated filtration through coffee filters, and stored at 4 °C until purification. Triculamin was purified by cation-exchange chromatography using an ÄKTA™ Start system equipped with a 5 mL HiTrap SP XL column (Cytiva). Six chromatographic runs were performed, each processing 2 L of culture supernatant at a flow rate of 5 mL $min^{-1}$, using the short program described below. Fractions were assessed for antimicrobial activity against *Mycobacterium smegmatis*. Fractions 5-8 from each run were pooled and desalted by dialysis against deionized water using a 500 Da molecular weight cutoff membrane.

Fractions 5–8 obtained from the ÄKTA cation-exchange purification were combined, lyophilized, and resuspended in 1 mL $H_2O$ containing 0.1% TFA. The combined sample was subjected to preparative HPLC. The preparative HPLC purification was performed using a Luna Omega 5 µm Polar C18 100 Å column (250 × 10 mm) with water containing 0.1% TFA as mobile phase A and acetonitrile containing 0.1% TFA as mobile phase B, at a flow rate of 4 mL/min. The gradient was held/increased from 5% B to 35% B over 25 min, followed by an increase to 100% B at 26 min, which was maintained until 35 min. The column was then returned to 0% B at 36 min and re-equilibrated until 45 min.

Baseline separation of triculamin A, B, and C could not be achieved in a single chromatographic run. Instead, the individual variants were isolated through repeated preparative HPLC runs, during which partially resolved fractions were pooled and reinjected until sufficient purity was obtained. Collected fractions corresponding to individual peaks were lyophilized and analyzed by HPLC-DAD and HPLC-MS. Purified fractions were lyophilized and weighed. The purified compound was used to make a dilution series and was run on UHPLC-DAD and HPLC-MS to generate a standard curve of triculamin A based on both MS signal (Fig. S2) and UV absorbance (Fig. S1).

#### 1.3 Expression and purification of triculamin variants from heterologous producer *Streptomyces albidoflavus* J1074

##### 1.3.1 Plasmid Construction

The overall cloning workflow is illustrated in Fig S3. Site-directed mutagenesis of the *pL99-ACDT* plasmid (12.2 kb, 66% GC) was performed using PCR amplification and Gibson Assembly. Due to the plasmid size and high GC content, mutations were introduced by amplifying two overlapping linear fragments using *pL99-ACDT* as a template. Each mutation was incorporated into one variable fragment, which was assembled with a common constant fragment to regenerate the full-length plasmid.

Primers were designed with 18–24 bp overhangs homologous to the vector termini to facilitate Gibson Assembly (see Table S5). For example, truncation mutants were generated by PCR amplification of a constant fragment using primers *trunc_cons_F* and *cons_R*, and variable fragments containing the desired deletions were amplified with *cons_F* and mutation-specific reverse primers (*del3_R*, *del6_R*, etc.). Point mutations in the core sequence were constructed similarly using primers such as *core_cons_F*/*cons_R* for the constant region and *cons_F*/*S1A_R* for the variable region.

PCRs were performed with Phusion™ Plus DNA Polymerase (Thermo Scientific) using the GC enhancer according to the manufacturer's protocol. Optimal annealing temperatures for each fragment were determined by gradient PCR (58–72 °C). Amplification products were verified by agarose gel electrophoresis and purified using the Monarch® DNA Gel Extraction Kit (NEB).

Purified fragments were assembled using NEBbuilder, and the resulting plasmids were transformed into chemically competent *E. coli* DH5α. Transformants were plated on LB agar containing apramycin (50 µg $mL^{-1}$) and incubated overnight at 37 °C. Colonies were screened by colony PCR (DreamTaq™ Green PCR Master Mix, Thermo Scientific), and positive clones were verified by whole-plasmid sequencing (Eurofins).

##### 1.3.2 Heterologous expression in *S. albidoflavus* J1074

Heterologous expression was performed in *S. albidoflavus* J1074 following the conjugation protocol described by Tong *et al.* (Crispr Best artikel). Briefly, purified plasmids carrying the different mutations in the *tri* biosynthetic gene cluster (BGC) were first transformed into electrocompetent *E. coli* ET12567. The resulting *E. coli* donor strains were co-cultivated with heat-shocked spores of *S. albidoflavus* J1074 on MS agar to enable interspecies conjugation. Prior to sporulation, nalidixic acid and apramycin (50 µg $mL^{-1}$) were added for negative selection against *E. coli*. Single exconjugant colonies were transferred to ISP2 agar containing apramycin (50 µg $mL^{-1}$) and subsequently spread onto MS agar plates supplemented with apramycin for sporulation. Plates were incubated at 28 °C for 3–5 days until full sporulation was achieved.

For expression of all mutations/truncation variants were cultivated in technical triplicates in 10 mL cultures. For all variants, the 3 cultures started from the same starter culture. For each variant, spores were harvested from two well-sporulated MS plates using 5 mL YEME medium without sucrose (YEME-WS) supplemented with apramycin (50 ug/mL) per plate. The resulting 10 mL of spore suspension was incubated 28 °C and 180 rpm for 24 h. The following morning 1 mL of the starter culture was used to inoculate 10 mL YEME-WS supplemented with apramycin (50 ug/mL). Cultures were grown in Erlenmeyer flasks at 28 °C and 180 rpm for 24 h, after which expression was induced by adding ε-caprolactam (0.5% w/v). Fermentations were continued for 5 days under the same conditions. The lysate was clarified by centrifugation (4000 rpm, 15 min) and stored at -20 °C until further analysis.

#### 1.4 Expression and purification of triculamin variants from *Burkholderia sp.* FERM BP-3421 Δfr9

##### 1.4.1 Plasmid construction

The plasmid generated for heterologous expression of the triBGC in *Burkholderia sp.* FERM BP-3421 was based on the two previously cloned plasmids: pL99-triACDT and pHNF-008-palABCD (ref. 1). Primers pal_F and pal_R were used with pHNF-008-palABCD for a PCR reaction according to previously described method. The PCR reaction was used for a gel-electrophoresis followed by gel-extraction of the correct size fragment. Primer's tri_F and tri_R were used with pL99-triACDT for a PCR reaction according to previously described method. The PCR reaction was used for a gel-electrophoresis followed by gel-extraction of the correct size fragment. The purified fragments were assembled using NEBBuilder and transformed into *E. coli* DH5a. The integrity of the plasmid was confirmed by whole plasmid sequencing. The workflow is illustrated in Fig S3.

##### 1.4.2 Expression of triculamin *Burkholderia sp.* FERM BP-3421 *Δfr9*

Heterologous expression in *Burkholderia sp.* FERM BP-3421 Δfr9 was performed according to the method described by Fernandez *et al.* (ref. 2,3). Briefly, electrocompetent cells were transformed with the target plasmid and selected on LB agar containing kanamycin (0.5 mg/mL). A single colony was used to inoculate 10 mL LB medium supplemented with kanamycin (0.5 mg/mL) and cultivated in a baffled Erlenmeyer flask at 28 °C and 180 rpm for 24h.

Due to interference of high kanamycin concentrations with downstream purification, the antibiotic concentration in the production medium was reduced to 50 µg/mL. The inoculum size was increased to a starting $OD_{600}$ of 0.1 (instead of 0.01). Production of the lasso peptide was carried out in 2S4G medium supplemented with kanamycin (50 µg/mL) and L-arabinose (100 mM), followed by incubation for 7 days. Cells and supernatants were collected by centrifugation (12,000 × g, 45 min), and triculamin was detected in the supernatant.

#### 1.5 Expression of triculamin in *E. coli* BL21(DE3)

For heterologous expression of triculamin in *Escherichia coli*, the same plasmid used for Burkholderia sp. FERM BP-3421 was employed. Chemically competent *E. coli* BL21(DE3) cells were transformed with sequence-verified plasmids for expression. Single colonies were used to inoculate Erlenmeyer flasks containing 10 mL LB broth supplemented with kanamycin (50 µg/mL), and cultures were incubated at 37 °C and 140 rpm for 18 h.

Overnight cultures were used to inoculate small-scale production flasks following the same workflow as described for *S. albidoflavus* and *Burkholderia* sp. FERM BP-3421. The following morning, three flasks containing 10 mL S4SG medium supplemented with kanamycin (50 µg/mL) were inoculated 1:100 with the overnight culture. In parallel, three flasks containing 10 mL LB medium supplemented with kanamycin (50 µg/mL) were inoculated in the same manner.

Cultures were incubated at 37 °C and 180 rpm, and $OD_{600}$ was monitored until reaching 0.4–0.8 (only measurable for LB cultures, as the presence of calcium carbonate in S4SG interferes with accurate $OD_{600}$ measurements). At this point, all six cultures were induced with L-arabinose (final concentration 0.5 mM) and incubated at 25 °C and 140 rpm for 18 h. After this a sample (2 mL) was taken from the flasks and the rest incubated an additional 2 days. Fermentations broths were stored at -20C, until further use.

#### 1.6 Cationic exchange purification of samples

All samples for all experiments were purified following the same method described here before any further analysis (UHPLC-DAD and HPLC-MS). Fermentation broths were purified using solid-phase extraction (SPE) with weak cation exchange cartridges (Strata-X-CW, 33 µm polymeric weak cation, 30 mg, Phenomenex). Samples were prepared by transferring 1200 µL fermentation broth to a 1.5 mL Eppendorf tube and heating at 95 °C for 10 min. After heating, samples were clarified (14,000 rpm, 10 min), and the supernatant was collected.

SPE columns were conditioned with 1 mL conditioning buffer 1 followed by 1 mL conditioning buffer 2. Subsequently, 1 mL of clarified sample was loaded. Columns were then washed with 1 mL wash buffer 1 and 1 mL wash buffer 2, followed by elution with 0.5 mL elution buffer, and the eluate was collected. Columns were reconditioned with 1 mL reconditioning buffer 1 and 1 mL reconditioning buffer 2 before reused. Each sample was processed in triplicate, and each column was reused three times (for the same mutation) before disposal. Eluates were collected in microcentrifuge tubes, frozen, and stored at –20 °C until further use.

#### 1.7 UHPLC analysis

Samples were analysed by UHPLC using a Luna® Omega Polar C18 column (100 × 2.1 mm, 1.6 µm particle size). Mobile phase A consisted of MQ water with 0.1% TFA, while mobile phase B consisted of acetonitrile with 0.1% TFA. The chromatographic gradient was run as follows: 0–0.6 min, 0–12% B; 0.6–5.0 min, 12–29% B; 5.0–5.5 min, 29–45% B; 5.5–6.0 min, 45–95% B; 6.0–10.5 min, 95% B; 10.5–11.0 min, 95–0% B; and 11.0–16.0 min, 0% B for column re-equilibration. Chromatographic data were analysed using the UV signal at 210.4 nm in OpenLab CDS software.

A standard curve was generated using this method and column under identical conditions, and the same setup was applied for analysis of all samples.

#### 1.8 HPLC–MS and MS/MS analysis

High-performance liquid chromatography coupled with tandem mass spectrometry (HPLC–MS/MS) was performed using an Orbitrap Exploris 120 mass spectrometer (Thermo Fisher Scientific) coupled to a Thermo Scientific Vanquish SII LC system. Chromatographic separation was achieved on a Luna Omega 1.6 µm Polar C18 100 Å, LC Column 100 x 1 mm. Mobile phase A consisted of $H_2O$ containing 0.1% formic acid, and mobile phase B consisted of acetonitrile containing 0.1% formic acid. Samples (10 µL) were injected following filtration through 0.22 µm membranes. The column temperature was maintained at 30 °C. The column was equilibrated for 5 min in 100% water before injection. Separation was performed using a water–acetonitrile gradient at a flow rate of 0.5 mL/min. After injection, the method was held at 100% water for 1 min, followed by an increase to 10% acetonitrile at 6.5 min, 30% acetonitrile at 12 min, 45% acetonitrile at 12.5 min, and 95% acetonitrile at 12.6 min. The run was stopped at 15.5 min.

The eluate from 2.5–12 min was directed into the mass spectrometer operating in positive electrospray ionization ($ESI^+$) mode. Source parameters were set as follows: spray voltage 3.5 kV, sheath gas 40 arbitrary units (AU), auxiliary gas 8 AU, sweep gas 1 AU, ion transfer tube temperature 300 °C, and vaporizer temperature 320 °C. Data-dependent collision-induced dissociation (CID) was employed for MS/MS acquisition at an Orbitrap resolution of 120,000 over a scan range of *m/z* 200–1000. The RF lens was set to 70%.

Raw data were processed using FreeStyle 1.8 for chromatographic inspection, spectral deconvolution, and MS/MS analysis. Some MSMS spectra (sFig. 4-12) were annotated using interactive peptide spectral annotator (ref. 4). Theoretical monoisotopic masses of triculamin B (tri-B, [M] = 1864.08205 Da) and its acetylated derivative (tri-B-Ac, [M] = 1906.09262 Da) were used for compound identification. These values correspond to dehydration of the linear tri-B precursor (1882.09735 Da) and subsequent acetylation (+42.01057 Da) for tri-B-Ac. Extracted ion signals at *m/z* 622.7022 ($[M+3H]^{3+}$) and *m/z* 636.371 ($[M+3H]^{3+}$) were used for comparison with verified standards and previously reported data (Merrild et al., 2025).

For quantification, deconvoluted spectra were extracted over a charge state range of 2–5, requiring detection of at least three charge states. Peak areas corresponding to theoretical masses within ±5 ppm were integrated into FreeStyle 1.8 and visualized using OriginPro 2018. For mutant variants, theoretical masses were adjusted accordingly (e.g., K2A: tri-B = 1807.02420 Da; tri-B-Ac = 1849.03477 Da).

##### 1.9 Statistics

Triculamin production values are reported as mean ± standard deviation (SD) from three fermentation replicates unless otherwise stated. Statistical comparisons were performed using two-sided Welch's t-tests against the positive control, defined as the non-mutated triculamin BGC expressed in *S. albidoflavus*. Welch's t-tests were used to allow comparisons without assuming equal variances between groups. P values were not adjusted for multiple comparisons.

Significance was annotated as ns, not significant; * $p < 0.05$; * $p < 0.01$; ** $p < 0.001$.

Concentration estimates were calculated using an external calibration curve generated with purified triculamin A. For MS-based total triculamin estimates, the summed MS signal from all detected triculamin variants was converted using the triculamin A calibration curve. These values should therefore be interpreted as approximate triculamin A-equivalent total triculamin concentrations, because the calculation assumes similar MS response factors across triculamin variants.

**Table S 1.** Overview of follower sequence truncations

| Deletion | Amino Acid Sequence |
|---|---|
| *TriA* | **MSKKSKPGDGIRGKGVRGR**FMFMDETPEAEATTSIRDSLADLAETLESEARK |
| *TriA_del3* | **MSKKSKPGDGIRGKGVRGR**FMFMDETPEAEATTSIRDSLADLAETLESE |
| *TriA_del6* | **MSKKSKPGDGIRGKGVRGR**FMFMDETPEAEATTSIRDSLADLAETL |
| *TriA_del9* | **MSKKSKPGDGIRGKGVRGR**FMFMDETPEAEATTSIRDSLADLA |
| *TriA_del12* | **MSKKSKPGDGIRGKGVRGR**FMFMDETPEAEATTSIRDSLA |
| *TriA_del15* | **MSKKSKPGDGIRGKGVRGR**FMFMDETPEAEATTSIRD |
| *TriA_del18* | **MSKKSKPGDGIRGKGVRGR**FMFMDETPEAEATTS |
| *TriA_del21* | **MSKKSKPGDGIRGKGVRGR**FMFMDETPEAEA |
| *TriA_del24* | **MSKKSKPGDGIRGKGVRGR**FMFMDETPE |
| *TriA_del27* | **MSKKSKPGDGIRGKGVRGR**FMFMDE |
| *TriA_del30* | **MSKKSKPGDGIRGKGVRGR**FMF |
| *TriA_del33* | **MSKKSKPGDGIRGKGVRGR** |

**Table S 2.** Overview of core and follower sequence mutations

| Mutation | Amino Acid Sequence |
|---|---|
| S1A | **MAKKSKPGDGIRGKGVRGR**FMFMDETPEAEATTSIRDSLADLAETLESEARK |

| | |
|---|---|
| K2T | **MSTKSKPGDGIRGKGVRGR**FMFMDETPEAEATTSIRDSLADLAETLESEARK |
| K3A | **MSKASKPGDGIRGKGVRGR**FMFMDETPEAEATTSIRDSLADLAETLESEARK |
| K3R | **MSKRSKPGDGIRGKGVRGR**FMFMDETPEAEATTSIRDSLADLAETLESEARK |
| P6A | **MSKKSKAGDGIRGKGVRGR**FMFMDETPEAEATTSIRDSLADLAETLESEARK |
| R11F | **MSKKSKPGDGIFGKGVRGR**FMFMDETPEAEATTSIRDSLADLAETLESEARK |
| V15A | **MSKKSKPGDGIRGKGARGR**FMFMDETPEAEATTSIRDSLADLAETLESEARK |
| R16A | **MSKKSKPGDGIRGKGVAGR**FMFMDETPEAEATTSIRDSLADLAETLESEARK |
| E29A | **MSKKSKPGDGIRGKGVRGR**FMFMDETPEAAATTSIRDSLADLAETLESAARK |
| L38S | **MSKKSKPGDGIRGKGVRGR**FMFMDETPEAEATTSIRDSSADLAETLESEARK |
| L41S | **MSKKSKPGDGIRGKGVRGR**FMFMDETPEAEATTSIRDSLADSAETLESEARK |
| L45S | **MSKKSKPGDGIRGKGVRGR**FMFMDETPEAEATTSIRDSLADLAETSESEARK |
| L38S, L41S, L45S | **MSKKSKPGDGIRGKGVRGR**FMFMDETPEAEATTSIRDSSADSAETSESEARK |
| L38S, L45S | **MSKKSKPGDGIRGKGVRGR**FMFMDETPEAEATTSIRDSSADLAETSESEARK |

**Table S 3.** List of media recipes.

| Name | Composition |
|---|---|
| LB Broth | Tryptone 10.0 g/L, yeast extract 5.0 g/L, NaCl 10.0 g/L |
| LB Agar | Tryptone 10.0 g/L, yeast extract 5.0 g/L, NaCl 10.0 g/L, agar 15.0 g/L |
| ISP2 Broth | Yeast extract 4.0 g/L, malt extract 10.0 g/L, glucose 4.0 g/L, pH 7.4 |
| ISP2 Agar | Yeast extract 4.0 g/L, malt extract 10.0 g/L, glucose 4.0 g/L, agar 15 g/L, pH 7.4 |
| MS Agar | D-mannitol 20.0 g/L, fat-reduced soy flour 20.0 g/L, agar 20.0 g/L, pH 8 After autoclavation add pre-autoclaved $MgCl_2$ (final concentration 10 mM) |
| 2xYT | Tryptone 16.0 g/L, yeast extract 10.0 g/L, NaCl 5 g/L |
| YEME Media without sucrose (YEME-WS) | Yeast extract 3.0 g/L, malt extract 3.0 g/L, peptone 5.0 g/L, glucose 10.0 g/L After autoclavation add pre-autoclaved $MgCl_2$ to final concentration 5 mM |
| Triculamin production media (TPM) | Glucose 22 g/L, yeast extract 2.5 g/L, $NH_4NO_3$ 4 g/L, $CaCO_3$ 2 g/L, NaCl 2 g/L |
| 2S4G media | Glycerol 40 g/L, soy peptone 20 g/L, ammonium sulfate 2 g/L, magnesium sulfate 0.06 g/L, calcium carbonate 2 g/L |

**Table S 4.** List of buffer concentrations.

| Name | Composition |
|---|---|
| SPE conditioning buffer 1 | 100% methanol |
| SPE conditioning buffer 2 /Wash 1 | 100 mM $NH_4COOH$, pH 10 |
| SPE wash 2 | 70% MeOH, 30% MQ water |
| SPE elution | 60% MeOH, 35% MQ water, 5% FA |
| Reconditioning buffer 1 | 50 mM NaOH |
| Reconditioning buffer 2 | 50 mM HCL |

**Table S 5.** List of primers.

| Name | Sequence (5'-3') |
|---|---|
| cons_F | GATGTACTCGGCGAGGTCGTTG |
| cons_R | CAACGACCTCGCCGAGTACATC |
| trunc_Cons_F | TAACCAGAGGCAGGTGACGG |
| core_cons_F | GAAGAGCAAGCCGGGCGAC |
| S1A_R | GTCGCCCGGCTTGCTCTTCTTCGCCATATGCGTACCTCCGTTGC |
| K2T_R | GTCGCCCGGCTTGCTCTTCGTCGACATATGCGTACCTCCGTTG |
| K2A_R | GTCGCCCGGCTTGCTCTTCGCCGACATATGCGTACCTCCGTTG |
| Leu_cons_F | GGAAAGCGAAGCCCGCAAG |
| L38S_R | CTTGCGGGCTTCGCTTTCCAGCGTCTCGGCCAGGTCGGCCGAGGAATCCCGGATCGAGG |
| L41S_R | CTTGCGGGCTTCGCTTTCCAGCGTCTCGGCCGAGTCGGCCAGGGAATCCCG |
| L45S_R | CTTGCGGGCTTCGCTTTCCGACGTCTCGGCCAGGTCGG |
| L38S, L41S, L45S R | CTTGCGGGCTTCGCTTTCCGACGTCTCGGCCGAGTCGGCCGAGGAATCCCGGATCGAGG |
| L41S, L45S R | CTTGCGGGCTTCGCTTTCCGACGTCTCGGCCGAGTCGGCCAGGGAATCCCG |
| glu_F | GGCCACCACCTCGATCCGGG |
| del3_R | CCGTCACCTGCCTCTGGTTATTCGCTTTCCAGCGTCTC |
| del6_R | CCGTCACCTGCCTCTGGTTACAGCGTCTCGGCCAG |
| del9_R | CCGTCACCTGCCTCTGGTTAGGCCAGGTCGGCCAGG |

| | |
|---|---|
| del12_R | CCGTCACCTGCCTCTGGTTAGGCCAGGGAATCCCGGATC |
| del15_R | CCGTCACCTGCCTCTGGTTAATCCCGGATCGAGGTGGTG |
| del18_R | CCGTCACCTGCCTCTGGTTACGAGGTGGTGGCCTCCG |
| del21_R | CCGTCACCTGCCTCTGGTTAGGCCTCCGCCTCCG |
| del24_R | CCGTCACCTGCCTCTGGTTACTCCGGGGTCTCGTCCATG |
| del27_R | CCGTCACCTGCCTCTGGTTACTCGTCCATGAACATGAAGCGTC |
| del30_R | CCGTCACCTGCCTCTGGTTAGAACATGAAGCGTCCCCGG |
| del33_R | CCGTCACCTGCCTCTGGTTAGCGTTCCCGGACGCC |
| Colony_PCR_F | CGTCGACCTGCAGGCATG |
| Colony_PCR_R | GAGCAGCCACAACAGTCC |
| pal_F: | CACGCCGTGAATGACGGCCCGTCAGATC |
| pal_R: | TCTTCGACATTTTAATCTTCCTAAAAGGTACCCGGG |
| tri_F: | GAAGATTAAAATGTCGAAGAAGAGCAAGCCGG |
| tri_R: | GGGCCGTCATTCACGGCGTGACGGTGG |

### 2 Supplementary results

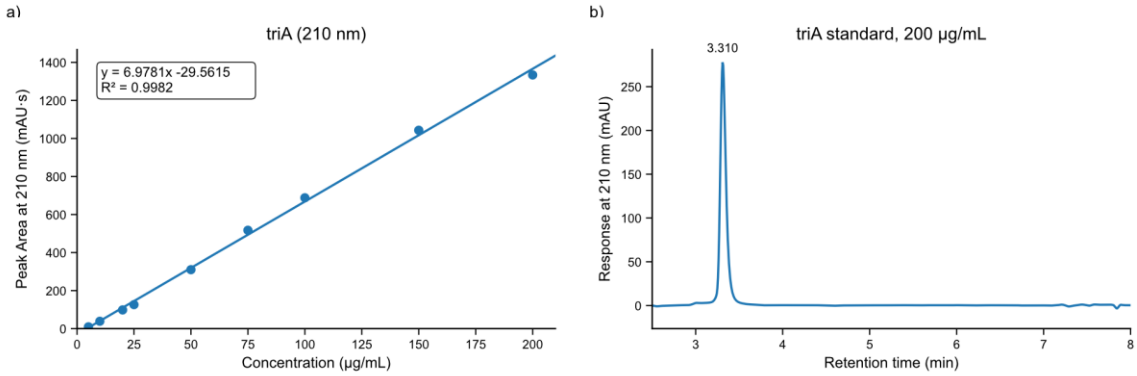


**Figure S1.** Standard curve for purified Triculamin A. a) UHPLC standard curve generated using purified Triculamin A isolated from *S. triculaminicus*. Peak area at 210 nm was measured across a concentration series and used to generate a linear regression model. The regression equation and coefficient of determination are shown on the plot. b) Representative HPLC-UV chromatogram of the 200 µg/mL triA standard measured at 210 nm.

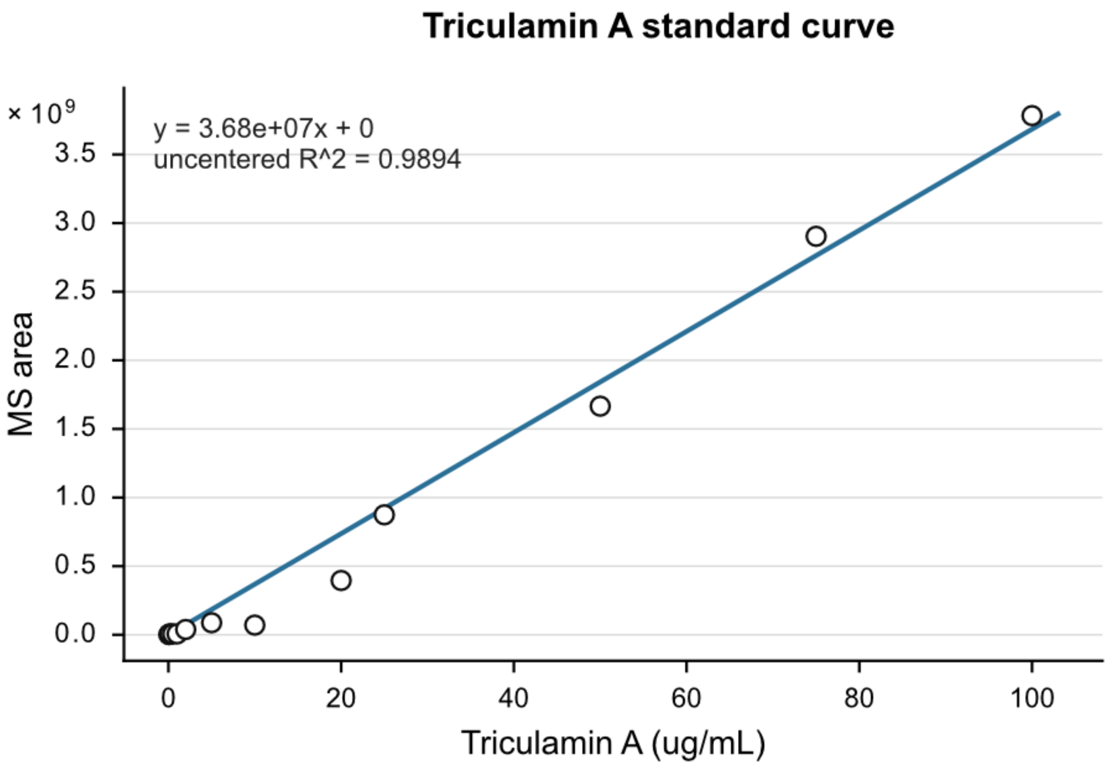


**Figure S2.** Triculamin A MS standard curve. Standard curve generated from triculamin A standards analyzed on the same MS system using 2 μL injections. The integrated MS area was plotted against triculamin A concentration and fitted by linear regression. The resulting equation was used to estimate triculamin concentrations in fermentation broth samples. Values were calculated from the deconvoluted MS signal of detected triculamin variants.

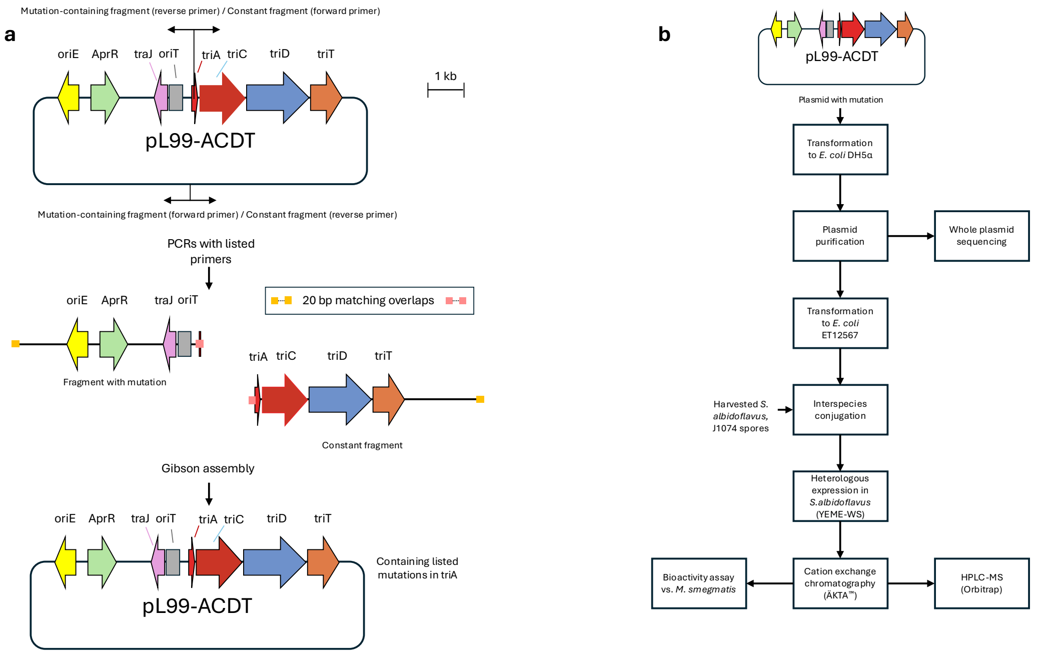


**Figure S3.** Cloning overview: Generation and screening of triA mutant plasmids. a) Schematic overview of triA mutagenesis in pL99-ACDT. Mutation-containing and constant PCR fragments were generated with matching 20 bp overlaps and assembled by Gibson assembly to produce pL99-ACDT variants containing the listed triA mutations. b) Workflow for transformation, plasmid validation, transfer into *S. albidoflavus*, heterologous expression, purification, LC-MS analysis, and bioactivity testing against *M. smegmatis*.

##### ***S. triculaminicus*** **produced triculamin A**

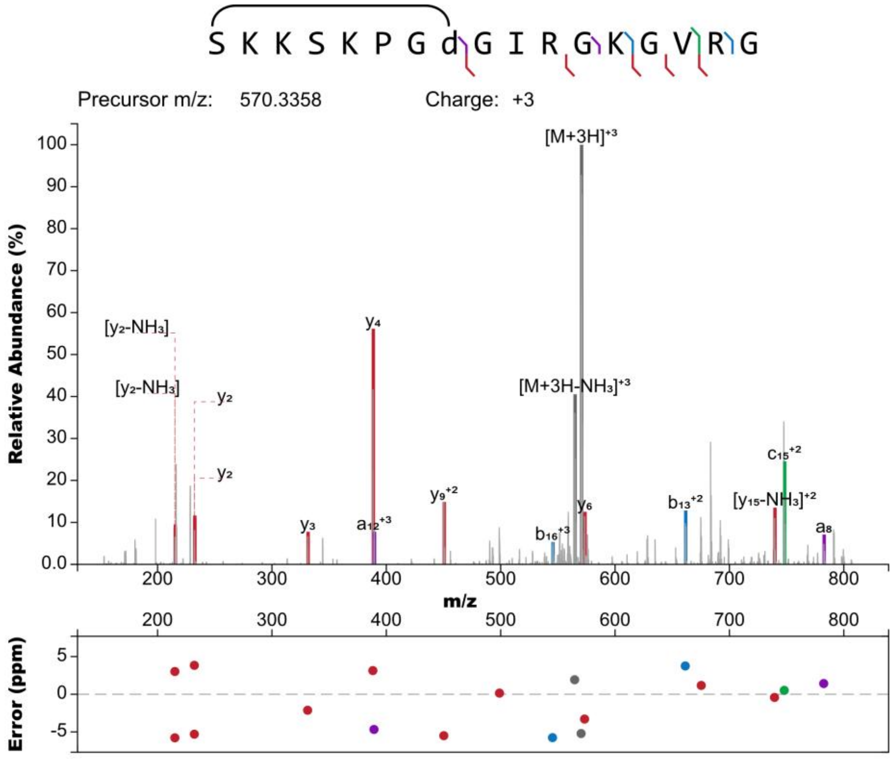


**Figure S4.** *S. triculaminicus* produced triculamin A MS$^2$ spectrum. Isopeptide bond between the N-terminus and aspartate is shown.

**_S. triculaminicus_ produced triculamin A-Ac**

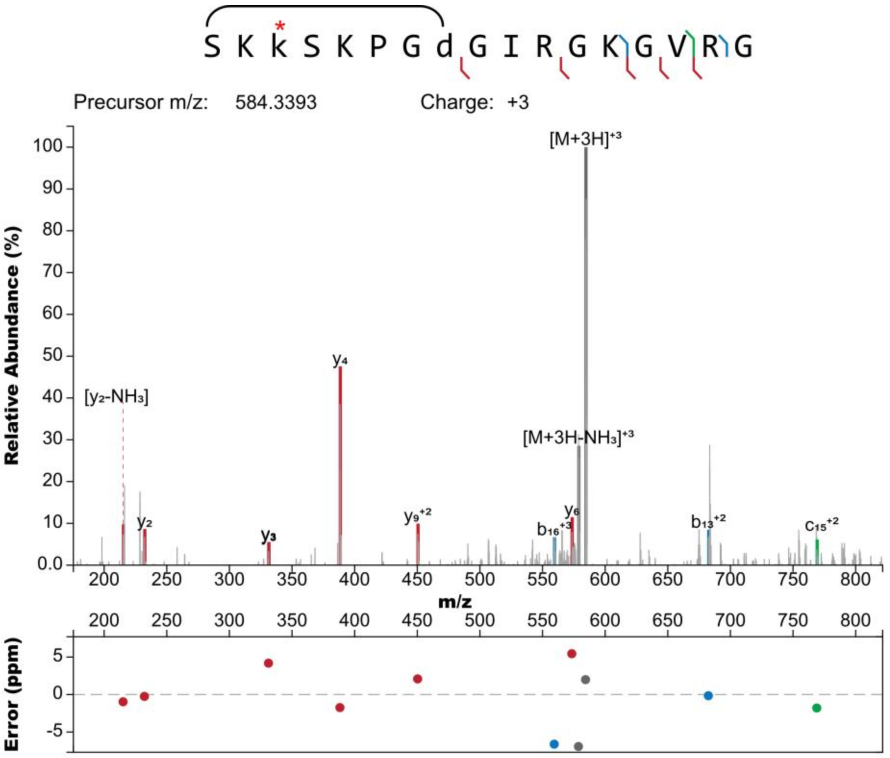


**Figure S5.** *S. triculaminicus* produced triculamin A-Ac $MS^2$ spectrum. Isopeptide bond between the N-terminus and aspartate and lys-3 acetylation is shown.

***S. triculaminicus* produced triculamin B**

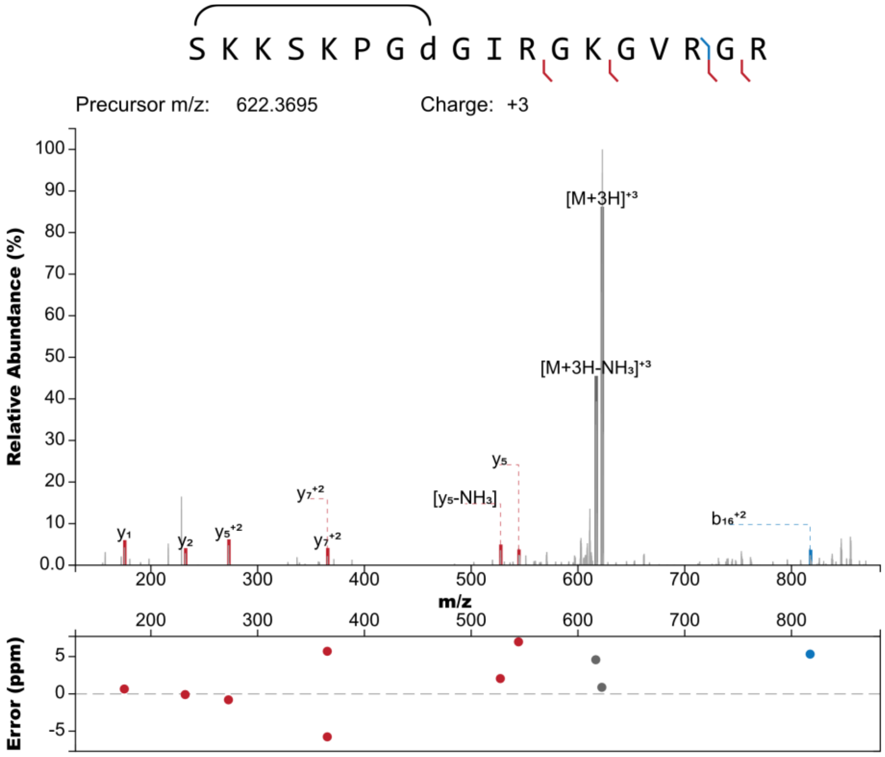


**Figure S6**. *S. triculaminicus* produced triculamin B MS$^2$ spectrum. Isopeptide bond between the N-terminus and aspartate is shown.

***S. triculaminicus* produced triculamin C**

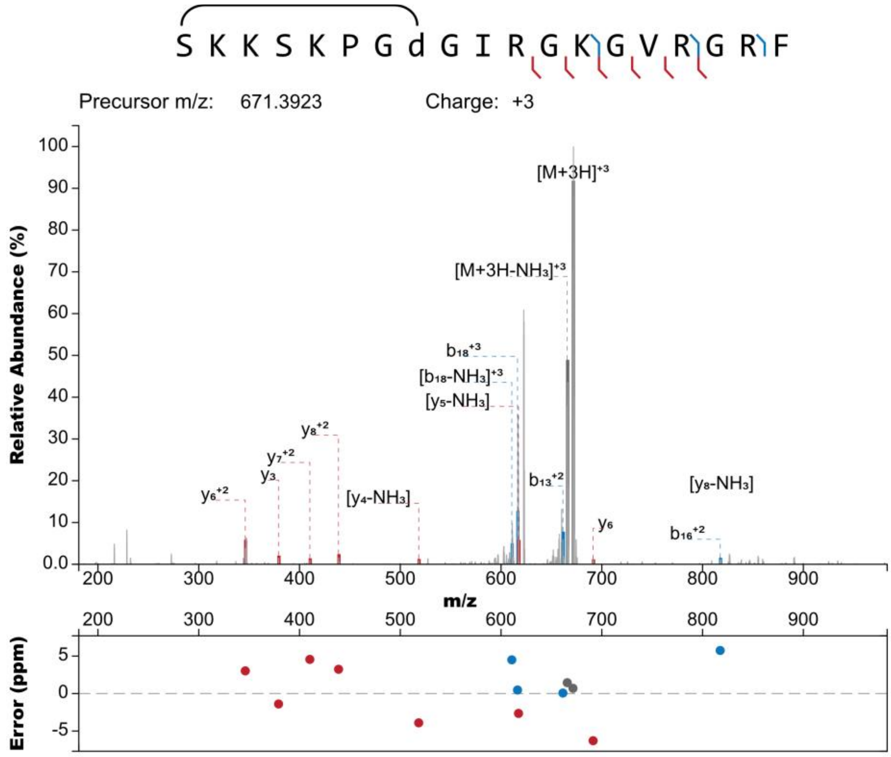


**Figure S7.** *S. triculaminicus* produced triculamin C MS[2] spectrum. Isopeptide bond between the N-terminus and aspartate is shown.

***S. triculaminicus* produced triculamin C-Ac**

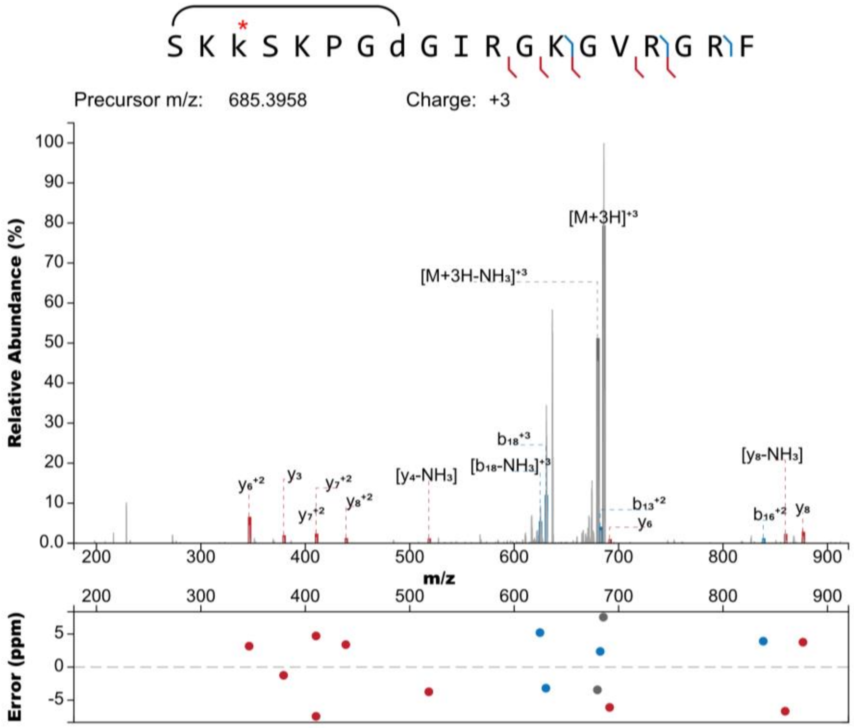


**Figure S8.** *S. triculaminicus* produced triculamin C-Ac $MS^2$ spectrum. Isopeptide bond between the N-terminus and aspartate and lys-3 acetylation is shown.

***Burkholderia* sp. FERM 3421 produced triculamin B**

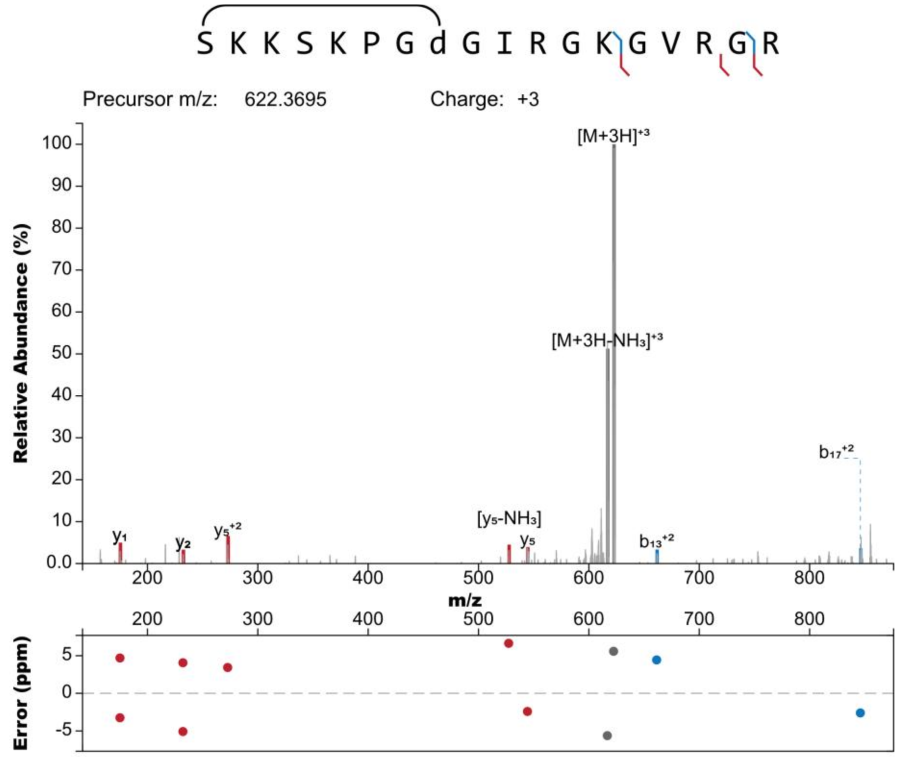


**Figure S9.** *Burkholderia sp.* FERM BP-3421 heterologously produced triculamin B $MS^2$ spectrum. Isopeptide bond between the N-terminus and aspartate is shown.

***Burkholderia* sp. FERM produced triculamin B-Ac**

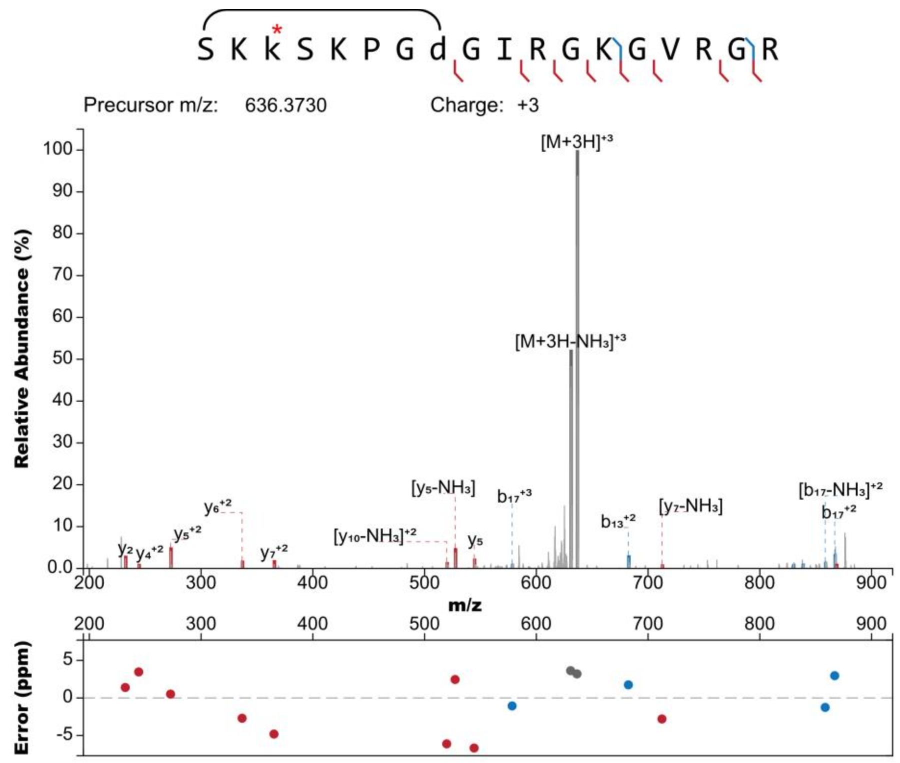


**Figure S10.** *Burkholderia sp.* FERM BP-3421 heterologously produced triculamin B-Ac $MS^2$ spectrum. Isopeptide bond between the N-terminus and aspartate and lys-3 acetylation are shown.

***Burkholderia* sp. FERM produced triculamin C**

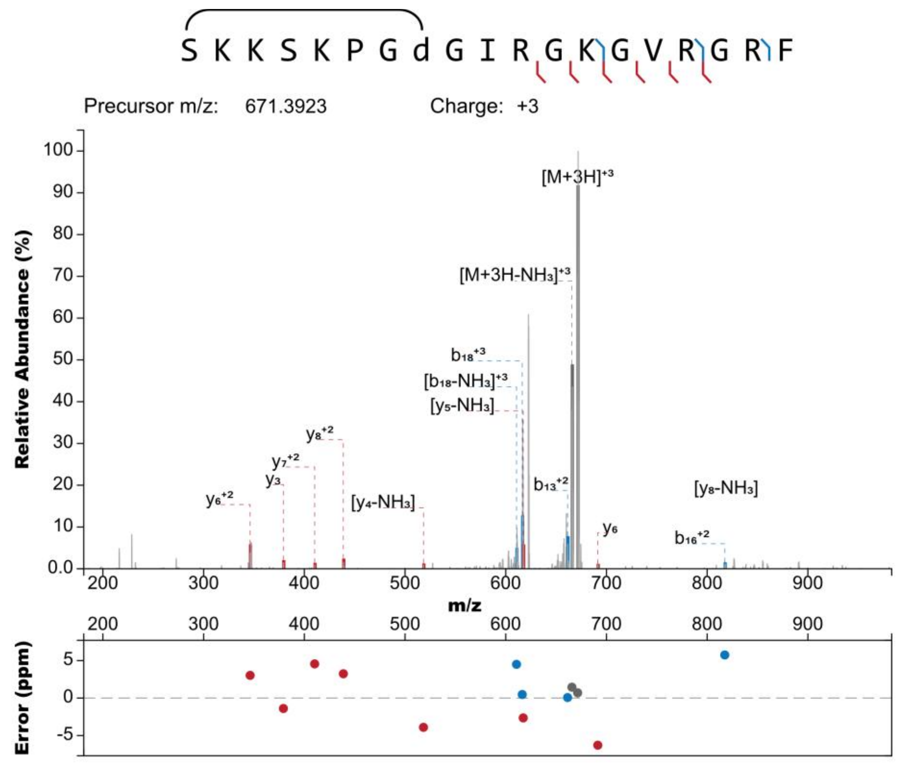


**Figure S11.** *Burkholderia sp.* FERM BP-3421 heterologously produced triculamin C $MS^2$ spectrum. Isopeptide bond between the N-terminus and aspartate is shown.

***Burkholderia* sp. FERM produced triculamin C-Ac**

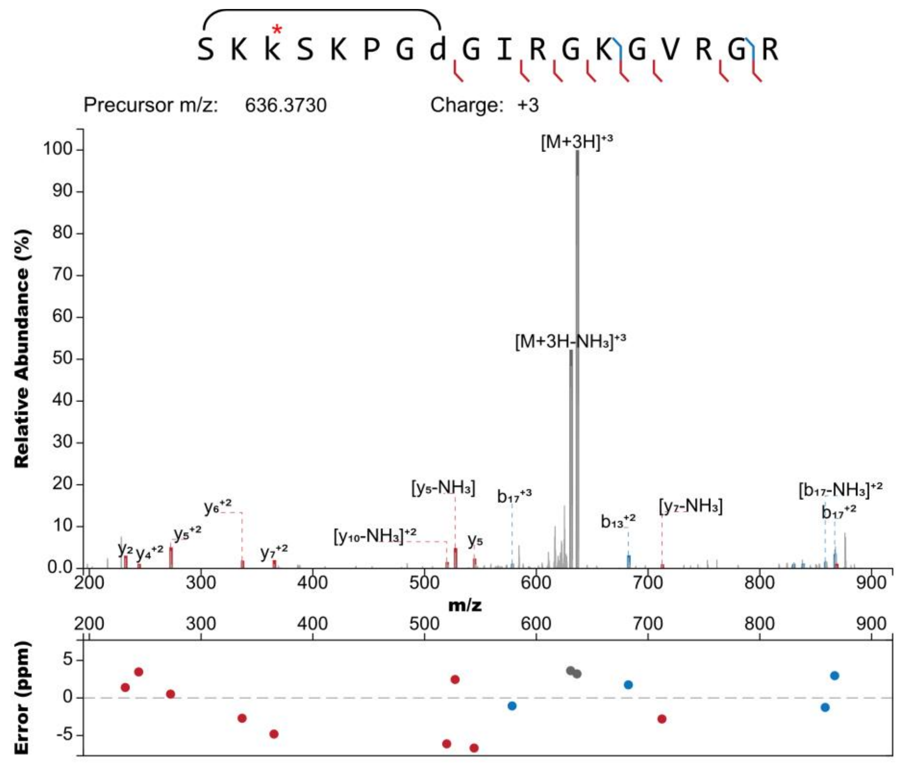


**Figure S12.** *Burkholderia sp.* FERM BP-3421 heterologously produced triculamin C-Ac MS[2] spectrum. Isopeptide bond between the N-terminus and aspartate and lys-3 acetylation are shown.

#### Constructed plasmid sequences

#### pHNF008-triACDTpalD used for heterologous expression in *Burkholderia sp.* FERM BP-3421

gccgcggggtcgtagaaccgtccgagcaggctgaagatcgtgctcttgccggccccggacggccctgcgatggccgtgaagctgtggcggggcacccggaacgacgccccccttgagaatctcgcgatcggtgtagcggaaccggaggtcgtgcagc
acgagcccggaggcggcgacggcgacggcgacgggctccttccggaactggtccgtctgcgggcggtccgtccggagccggtccgtttcgacgggcagatccaggacctcctggatgcgctgcagcgccgcgagcccggcttgcagtgcgggagcg
atgtcgaacagcccggccatcggtacggccaggtagttcacgtagaggaggaaggccaccagctcgccgacggccatgtccccggacgcgaccctggccaggccgacgacgatgaccacgatcagcgagccctgcaccgcgagttcgatggcgg
gtgagaccacggcggtcagacgggcggccttgacgttgtgaccgtagatgtcgtcggtgctacgggccagcgccgcggattcgcgcgcctcggcacggttgatgcgcacggtgcggatggcgctgaggacgcgctccagctcggcggcgagggagcc
ggtggcttcctgcgcggcctgggtctcgcggcggatccccttgagtacgagcgccatcaccgctccggacccgatgacgaccgccatgaccaccccggcgagtatcggatccagccagaacatcatgcccacgccgccgaccacgacgatggagt
tgacgatcgcgttcaccacgtcggtggcgagcacgtcgcgcagcagggtggtgtcggtggtggtgcgggagatcaggtcgcccagccggtgacggtcgtagacggccatgcgcaagcacagcaactggtcgaccatcgccctgcgcagccgctggac
gacaccctcgccggcctggtcgagccggaacgtcgaccaggcctgcaatgcggcttggcctgcgaagacggcggcgagcagccacaacagtccggcgacggaccgccccgccgtcgccgcctcgatggtccggttgacgatcagaggctgggcc
agcccgagtccggatccggccagggcgcagagcagagcgaacagcagccttcgccgactgccccgcgtccaggccagcagagtacggaaggtgacgcggctgactttcgaggtttcgtccatggctccgcaagggtgattagtcgatcagttcgtaa
ccgagttccatcgccgtggccagccattgctcgacctcgacgatgcgggtcgccaccatgggatccaggtctcctgacagcgcgcgggcgctgaggatcccgctctccagcagatacttcgccgacagttcggcgggatccacacggctgacgatgg
cccggctgaagatttgtttggcggtggggagatgcgccagggcacccgggggcatcagcttgatcacctgggatttcatccgccagtacgggtgcgggtagctcgggtcgtagcgccggtgcagcggtagtcgcttcgcccgttcgacgaactccgggtc
caggaaggggtcctcgttgcgcaccagaccggctgtgcgcagccgttggtgtggatagaggtcgctccacgcttccatgacggccagcgaatcatggtgccgttcgtggagccgtaccagcgaaccgatccaggacttcgtccatgcggtggcgattcc
tcggtacggctcggcgaggaagtcgtgcggtggattgctgcacagctccggccaggaacacgccagatacgtcatcgcccggcgtcgtcggccgaggaggggcgagagtacggcgagcgtctcgagtttcttcgtgggctgatactcgccgcggtgatc
gcgccagtagtccggcagtgcgcgcacgtttccgtggcgaaacaggtggggtgtcaggtactgcccgctgcccaggagttcgtcggcgccgtcgccggtgagcagcacgtccgcgcccgcctcgtgggcgagttccaccagaagccggttggcatcg
ggcagggcgtcgagacgtggcccttccgcatgccattccggaaccccgtcggtcgccgggagatcggtacgatccgcaggtcgcaccgcagaccgattccctcgagcaggttccgggcgacggggatgttggacttcccccggtcatcgatcatttc
gatggcgacagcggtgatccggcgctgatcaccgtgctcgaggatgaggttggccagaacggcgagggagtcgagcccgccggaggtggcgaccgcgacccccctcgctgttcccgacggcgcgccgcagcgaggcgaggaactccgtgcgcagat
ttcgcattccaggttcgccccggggccacgggggttccgtcgccatatccggaccgtcctcgaagggatgggccgggccgaactcgcccctgatggccggccaggtcaaccacagtgcgagtctgcgtcgggagatccgctcgtacggcatgccgtc
acctgcctctggttacttgcgggcttcgctttccagcgtctcggccaggtcggccagggaatcccggatcgaggtggtggcctccgcctccggggtctcgtccatgaacatgaagcgtccccggacgcccttgccgcgaataccgtcgcccggcttgctc
ttcttcgacattttaatcttcctaaaaggtacccggggatcctctagagtcgaccccaaaaaaacgggtatggagaaacagtagagagttgcgataaaaagcgtcaggtgggatccgctaatcttatggataaaaatgctatggcatagcaaagtgtgacg
ccgtgcaaataatcaatgtggacttttctgccgtgattatagacacttttgttacgcgtttttgtcatggctttggtcccgctttgttacagaatgcttttaataagcggggttaccggcttggttagcgagaagagccagtaaaagacgcagtgacggcaatgtc
tgatgcaatatggacaattggtttcttctctgaatggcgggagtatgaaaagtatggctgaagcgcaaaatgatcccctgctgccgggatactcgtttaatgcccatctggtggcgggtttaacgccgattgaggccaacggttatctcgatttttttatcgacc
gaccgctgggaatgaaaggttatattctcaatctcaccattcgcggtcaggggggtggtgaaaaatcagggacgagaatttgtttgccgaccgggtgatattttgctgttcccgccaggagagattcatcactacggtcgtcatccggaggctcgcgaatggt
atcaccagtgggtttactttcgtccgcgcgcctactggcatgaatggcttaactggccgtcaatatttgccaatacggggttctttcgcccggatgaagcgcaccagccgcatttcagcgacctgtttgggcaaatcattaacgccgggcaaggggaaggg
cgctattcggagctgctggcgataaatctgcttgagcaattgttactgcggcgcatggaagcgattaacgagtcgctccatccaccgatggataatcgggtacgcgaggcttgtcagtacatcagcgatcacctggcagacagcaattttgatatcgccag
cgtcgcacagcatgtttgcttgtcgccgtcgcgtctgtcacatcttttccgccagcagttagggattagcgtcttaagctggcgcgaggaccaacgtatcagccaggcgaagctgcttttgagcaccacccggatgcctatcgccaccgtcggtcgcaatg
ttggttttgacgatcaactctacttctcgcgggtatttaaaaaatgcaccggggccagcccgagcgagttccgtgccggttgtgaagaaaaagtgaatgatgtagccgtcaagttgtcataataaatcgatgcagggctagcggggccctgatatctggtgct
acgcctgaataagtgataataagcggatgaatggcagaaattcgaaagcaaattcgacccggtcgtcggttcagggcagggtcgttaaatagccgcttatgtctattgctggtttaccggtttattgactaccggaagcagtgtgaccgtgtgcttctcaaat
gcctgaggccagtttgctcaggctctccccgtggaggtaataattgacgatatgatcatttattctgcctcccagagcctgataaaaacggtgaatccgttagcgaggtgccgccggcttccattcaggtcgaggtggcccggctccatgcaccgcgacgc
aacgcggggaggcagacaaggtatagggcggcgaggcggctacagccgatagtctggaacagcgcacttacgggttgctgcgcaacccaagtgctaccggcgcggcagcgtgacccgtgtcggcggctccaacggctcgccatcgtccagaaaac
acggctcatcgggcatcggcaggcgctgctgcccgcgccgttcccattcctccgtttcggtcaaggctggcaggtctggttccatgcccggaatgccgggctggctgggcggctcctcgccggggccggtcggtagttgctgctcgcccggatacagggt
cgggatgcggcgcaggtcgccatgccccaacagcgattcgtcctggtcgtcgtgatcaaccaccacggcggcactgaacaccgacaggcgcaactggtcgcggggctggccccacgccacgcggtcattgaccacgtaggccgacacggtgccg
gggccgttgagcttcacgacggagatccagcgctcggccaccaagtccttgactgcgtattggaccgtccgcaaagaacgtccgatgagcttggaaagtgtcttctggctgaccaccacggcgttctggtggcccatctgcgccacgaggtgatgcagc
agcattgccgccgtgggtttcctcgcaataagcccggcccacgcctcatgcgctttgcgttccgtttgcacccagtgaccgggcttgttcttggcttgaatgccgatttctctggactgcgtggccatgcttatctccatgcggtaggggtgccgcacggttgc
ggcaccatgcgcaatcagctgcaacttttcggcagcgcgacaacaattatgcgttgcgtaaaagtggcagtcaattacagattttctttaacctacgcaatgagctattgcggggggtgccgcaatgagctgttgcgtaccccccttttttaagttgttgatttt

taagtctttcgcatttcgccctatatctagttctttggtgcccaaagaagggcacccctgcggggttcccccacgccttcggcgcggctccccctccggcaaaaagtggcccctccggggcttgttgatcgactgcgcggccttcggccttgcccaaggtg
gcgctgccccccttggaacccccgcactcgccgccgtgaggctcggggggcaggcgggcgggcttcgcccttcgactgcccccactcgcataggcttgggtcgttccaggcgcgtcaaggccaagccgctgcgcggtcgctgcgcgagccttgaccc
gccttccacttggtgtccaaccggcaagcgaagcgcgcaggccgcaggccggaggcttttccccagagaaaattaaaaaaattgatggggcaaggccgcaggccgcgcagttggagccggtgggtatgtggtcgaaggctgggtagccggtgggcaa
tccctgtggtcaagctcgtgggcaggcgcagcctgtccatcagcttgtccagcagggttgtccacgggccgagcgaagcgagccagccggtggccgctcgcggccatcgtccacatatccacgggctggcaagggagcgcagcgaccgcgcagggc
gaagcccggagagcaagcccgtagggcgccgcagccgccgtaggcggtcacgactttgcgaagcaaagtctagtgagtatactcaagcattgagtggcccgccggaggcaccgccttgcgctgcccccgtcgagccggttggacaccaaaaggga
ggggcaggcatggcggcatacgcgatcatgcgatgcaagaagctggcgaaaatgggcaacgtggcggccagtctcaagcacgcctaccgcgagcgcgagacgcccaacgctgacgccagcaggacgccagagaacgagcactgggcggccag
cagcaccgatgaagcgatgggccgactgcgcgagttgctgccagagaagcggcgcaaggacgctgtgttggcggtcgagtacgtcatgacggccagcccggaatggtggaagtcggccagccaagaacagcaggcggcgttcttcgagaaggcgca
caagtggctggcggacaagtacggggcggatcgcatcgtgacggccagcatccaccgtgacgaaaccagcccgcacatgaccgcgttcgtggtgccgctgacgcaggacggcaggctgtcggccaaggagttcatcggcaacaaagcgcagatga
cccgcgaccagaccacgtttgcggccgctgtggccgatctagggctgcaacggggcatcgagggcagcaaggcacgtcacacgcgcattcaggcgttctacgaggccctggagcggccaccagtgggccacgtcaccatcagcccgcaagcggtc
gagccacgcgcctatgcaccgcagggattggccgaaaagctgggaatctcaaagcgcgttgagacgccggaagccgtggccgaccggctgacaaaagcggttcggcaggggtatgagcctgccctacaggccgccgcaggagcgcgtgagatgcg
caagaaggccgatcaagcccaagagacggcccgagatcttcgggagcgcctgaagcccgttctggacgccctggggccgttgaatcgggatatgcaggccaaggccgccgcgatcatcaaggccgtgggcgaaaagctgctgacggaacagcggg
aagtccagcgccagaaacaggcccagcgccagcaggaacgcgggcgcgcacatttccccgaaaagtgccacctgacgtctaagaaaccattattatcatgacattaacctataaaaataggcgtatcacgaggccctttcgtcttcgaataaatacc
tgtgacggaagatcacttcgcagaataaataaatcctggtgtccctgttgataccgggaagccctgggccaacttttggcgaaaatgagacgttgatcggcacgtaagaggttccaactttcaccataatgaaataagatcactaccgggcgtatttttga
gttatcgagattttcaggagctaaggaagctaaaatggagaaaaaaatcactggatataccaccgttgatatatcccaatggcatcgtaaagaacattttgaggcatttcagtcagttgctcaatgtacctataaccagaccgttcagctggatattacggc
ctttttagcttttgccattctcaccggattcagtcgtcactcatggtgatttctcacttgataaccttatttttgacgaggggaaattaataggttgtattgatgttggacgagtcggaatcgcagaccgataccaggatcttgccatcctatggaactgcctcggt
gagttttctccttcattacagaaacggctttttcaaaaatatggtattgataatcctgatatgaataaattgcagtttcatttgatgctcgatgagtttttctaatcagaattggttaattggttgtaacactggcagagcattacgctgacttgacgggacggcggct
ttgttgaataaatcgaacttttgctgagttgaaggatcagatcacgcatcttcccgacaacgcagaccgttccgtggcaaagcaaaagttcaaaatcaccaactggtccacctacaacaaagctctcatcaaccgtggctccctcactttctggctggat
gatggggcgattcaggcctggtatgccgcggcagatgtctgcctgttcatccgcgtccagctcgttgagtttctccagaagcgttaatgtctggcttctgataaagcgggccatgttaagggcggtttttttcctgtttggtcactgatgcctccgtgtaaggggga
tttctgttcatgggggtaatgataccgatgaaacgagagaggatgctcacgatacgggttactgatgatgaacatgcccggttactggaacgttgtgagggtaaacaactggcggtatggatgcggcgggaccagagaaaaatcactcagggtcaatgcc
agcgcttcgttaatacagatgtaggtgttccacagggtagccagcagcatcctgcgatgcagatccggaacataatggtgcagggcgctgacttccgcgtttccagactttacgaaacacggaaaccgaagaccattcatgttgttgctcaggtcgcaga
cgttttgcagcagcagtcgcttcacgttcgctcgcgtatcggtgattcattctgctaaccagtaaggcaaccccgccagcctagccgggtcctcaacgacaggagcacgatcatgcgcacccgtggggccgccatgccggcgataatggcctgcttctc
gccgaaacgtttggtggcgggaccagtgacgaaggcttgagcgagggcgtgcaagattccgaataccgcaagcgacaggccgatcatcgtcgcgctccagcgaaagcggtcctcgccgaaaatgacccagagcgctgccggcacctgtcctacga
gttgcatgataaagaagacagtcataagtgcggcgacgatagtcatgccccgcgcccaccggaaggagctgactgggttgaaggctctcaagggcatcggtcgagatcccggtgcctaatgagtgagctaacttacattaattgcggaattcaagcact
agtttaattaatcagaagaactcgtcaagaaggcgatagaaggcgatgcgctgcgaatcgggagcggcgataccgtaaagcacgaggaagcggtcagcccattcgccgccaagctcttcagcaatatcacgggtagccaacgctatgtcctgatagc
ggtccgccacacccagccggccacagtcgatgaatccagaaaagcggccattttccaccatgatattcggcaagcaggcatcgccatgggtcacgacgagatcctcgccgtcgggcatccgcgccttgagcctggcgaacagttcggctggcgcga
gcccctgatgctcttcgtccagatcatcctgatcgacaagaccggcttccatccgagtacgtgctcgctcgatgcgatgtttcgcttggtggtcgaatgggcaggtagccggatcaagcgtatgcagccgccgcattgcatcagccatgatggatactttct
cggcaggagcaaggtgagatgacaggagatcctgccccggcacttcgcccaatagcagccagtcccttcccgcttcagtgacaacgtcgagcacagctgcgcaaggaacgcccgtcgtggccagccacgatagccgcgctgcctcgtcttggagtt
cattcagggcaccggacaggtcggtcttgacaaaaagaaccgggcgcccctgcgctgacagccggaacacggcggcatcagagcagccgattgtctgttgtgcccagtcatagccgaatagcctctccacccaagcggccggagaacctgcgtgc
aatccatcttgttcaatcatactcttcctttttcaaattattgaagcatttatcagggttattgtctcatgagcggatacatatttgaatgtatttagaaaaataaacaaataggggttccgcgcacatttccccgaaaagtgccacctgacgtctaagaaacca
ttattatcatgacattaacctataaaaataggcgtatcacgaggccctttcgtctcgcgcgtttcggtgatgacggtgaaaacctctgacacatgcagctcccggagacggtcacagcttgtctgtaagcggatgccgggagcagacaagcccgtcgcat
gcatctagagggatccgctgcagggcggccgctcaggtcattacatcggcgtgtccggttgtgcctggtagaactgggcctggctacggaacatacgtgcgtattttccgtcgcgtgcgagcagcgcatcatgcgagcctgtttcgacgattcggccctgct
cgagcaccacgatgcgatccacgaaccgcgtcagtgcgagtcgatggctgacgatcacggtgctccgccgtctggcgatacgcaggatcaatcgcatgatctccgcctcgaattcggggtccagcgcggatgtcggttcgtcgaaaaagagcaggtcg
gcctcgtcggcatgggccagcacgcgcgcgatggcgagtcgttgccattggccaccggagagatcgacgccatcggccatttcgcttgtcagtggctgcgtcaagcgcgtgctcagactgtccagacccaccgcgcgcaggtgggcccttgcccggct
gccatcgcgctcgtcgggcataccgagattcgcgatcacgtcgaatgggaagcgcgcgaaatcctgaaacagaccgacgacgcgcggcggcgccggctcggtcgcccaacgcagggtgccggcggtcggttggtagaggccgctcaacagcttga
gcaagctggtcttgcccgcgccgttttcaccgacgacggcgatcgactctcgcggtgcaatctccaggtcgacgtcgacgagcgcgggccgcgccgcgtcccggtaatgaaggctcacgccacggagggaaacgctgcttgccgccggcgcggcaa
caccatcgatgccgcctccgtcgggtgtcggctggcttgccctatcacgttgacgcgcggcaaggtgggccgccgtcaacgcgcgtagcgggcgcgcggcataggacacgcgcagtacgtcgccaacgttgtagatgattgccgccaggccgtctttc
aattgcaccaacgcgccgagcaggaaagcgagctcggcaatggtgaaacggcccgctgcacagcctgccacgacgacgtacaggggtatggccaggcagccgctcgcggcaatagaggtcccggcaagcttgagggcgttgcgtgtgcgcacgc
gccgcatcgaatcgagataggtcttgtaacgatgccgccagcccgcgagcaggtcctgctgcatccggtagacgcgcaggtccttcgcgtactccggctgcgtcagcacccgttcgacgatgcgcagcgcattgaaggtcccggcatggcgctcgcgt
acatcccagctctcgcgttccgcccgggctcggcagtacacggcgggtgtcatccctgccagcatcagcagcgggatccaccacgcggtgtgtgcggtaatgacgaggatcggcgtcatcatcacggtaccaaggcagacggcgtacagatgctgga
ccagatcgccgatctcgtcgctggccgcggccgcgaggagcgccgtctcgcgtagctgggcatcttcgtgaatgctcaggtctggataggttgagacgatcgcgttgacctggcccttcaatcgcactttcgccgcgtcctcgagcgtatcgacgacgaa
gctcgaaatggcgttcagcccgtcgtggcaaccgccaagcaggaagtagctgaataccagtgccatgacggccgcgacgccggcgttcgagctgaggcgaccgatcagttgcgcgttgatgtagagcaacagcgccgggatggcaccgaccacca
ggttggacgcgagcgcgagacacattgcgcgcggtgccgcgtgccacagcagcgacagcacttcgtatagattggcgacgatctgacgggccgtcattcacggcgtgacggtgggccgtacgtagatccacagctgatcgacttcgttgaatccggcg
gattcgtacagggcgttggccgccgtccggtcgcgttcgcccccgggggggttcgtcgtcgtcgacgtagagaatcgcttcagtggctcctagaccggtgagcgtgtcgagggcggagccgagtagtgcgcgtcccaggccctgccttcgcgcttcggggc
cgacctcgatccaccacaggacgccgacgccctggacgggttcgccgatgaccgcctcggccaccgtccggccgtctctgcgcacctgcagttgccgagcggtcgtccgctcgggctcggggacgatccggatgttgtccgcccgtggcaggccgg
ggcgcggtagggcggcgtgcatgtagcgccagaggcgttcgccggtgaacccggcgcgttccagtgccgcgcgggtcaccggccggtggccgatcggcagcgcctccagacccagggtgagcgcggtcgcgaactggaaggcttcgacggggcgg
ggcgccagctctgccagggcatgggcgatcagcgcctcggccacggccgcgttctcacggcagtgggcccacaggatcagcccggtgtcgtctttcggacgcagtgcgcaggagacgacgcccacgacgttgccgttgccgtcggcggtggtggcca
cgtgtacggtcggcgggtcgagttcggcccaccagttgtcgtcgacgggcgactcgccgcgcagggcctgcgcgagcatgtcgggtgtggcttccggctggcccgggagcctgtcggcgttgacaagggtcaggacggccgtccggtcggtcgcgcggt
acgggcggagcgttaggcgtgtcatggctaccacactggtcatcgggtcctcttctgtccaggacggccggttgctcgtcgcccgtaccggtcatcaggcggccgcgtgctgcgcgggcaggtggtcccgtatctcgtcgtagccgccggcggcgacca
cccgcccgccgtccagcacgacgacctggtcggcggaccggatcgtggacagccggtgggcgatgaccagcagcgtgcactccttggagatcagatgcagggtctgtgccaggagctgctcatgggcggcgtccaggtgcgaggtgggctcgtccag
caacaggatctccggccgggccagcagggcacgagcgatcgccacgcgctgttgttcgccaccggacaacgacgagccgtgttccccgacgggggtgtcgaggccgtcggcagccgcttcaccaaggcggacaagcccgccatctccactacc
cggtgcagccacggttcttcggcgtccggcgccgcgtaggtgaggttgtcgcgcagagtgccgtacatcagcggtgaatgctgctccaccaggccgatgcgggaccgccattcgcccgtggtgtgctcggtgtgggcatcgcgcccgtcgagggtgatg
ctcccg

#### Plasmid pL99-triACDT used for cloning and expression of precursor mutants:

tcaccgacccccgccgtacgtcaccgggatgacgtacggcgggggggagcgagttagtgcgaagtgggcccacttgcgagccgggcgatgtgccgggcggcccgctcctggcggtcgtcggcgtcgtcgtcctggtcgtcgtcctgctctcgccgtcg
gcgtgcagttgcttcctcgcggcgctgggcgagggcggcgagcatgtcggcgtacgcctcggccacctcccccgccgtgagcaccaccactgtgtcggccgcgtcggccagcgccaggacctcccgcacccgttcgcccacggccgccgaatcct
cgttgccgtccttgccttcggcggcccgggtcgcctcgaggtcgagggcgcggcgggtgaccgcgtgccatccgtcctcggtcacggcgaccccggcccgcagctccccgccgtcggcgtcggccgccaggagcagatcgaggtcgtcggcctcggt
gtcgccgccgtcgagcccgagcatctgccgcaggtagcgggtccattcgatggcccggcgtccccgggttgcccgctcgtactcgtgccagcgcgagaggttccactccagcgagccgaccccggcggcgtcgtcctcggtcatgccgccggtcagg
tccccgatccgtccgaggagttcgaacggggcgacgttcccgccggtcgccgtcttgaggtcggcgcgggcgagttcgagggcgggcgccttcccgtcctgggtcttggcgatgtactcggcgaggtcgttggcgtcgcgctcggtctccagccgcttga
agtcgacgccgtgccggtcgtcgggcgtgaaggcggggttgaccttgcgcagggcggcggtccacacggaccgccagtgcccctgccactcgtcgagcgcggcgccggtcggctcgaaggtggcgacgatctgcttcgcggaccgctccccctcggt
ccggccgccgaccaggacgatcgcgtggatgtgcgggtgccagccgttgatctgccccacggtgacttcggtcgcgcggatcatgccgacgtacccgatccggtctcggatgccctcgcggtcggcggcccggtgcccgtccttggcccggcgtccgg
cccacgtgccgcccgtgatcagtcgctggtaggcgcccggccgccgggggctgtccggcgtcttccgggtgccctggagggcgtccatgaggtccgcgagccggtccgtgtgcccatggcgggccgtgaaggtgaccaggtaggcggtccccccgcg
cttgatccactcgaccacggcggcggtgatctcctcggcccgcttgtgccggatcgtggcggcgcagaccgggcagagccagatccgcccgcaccgcatcaggcccaggaccacggacgttccggccgccgtctgggcgacgattacgccggagg
cagggtccatcagggcgcggccgcagcccttgcacgcggcgtccccgctgatccgccacagcgtccggcggcggctgtaccgggcggctttccgcagtcgggcagcgtcgctccgcgacgtgcttcctacttccgagaggctgtcgcctctcgggctc
tccccatccaccccgtccggagaaaccgcaggtcggaggggtgcgggaaactctgttgtttctttcccaaggtgttcgcttttgcctcgggcggcatctcgcgtcacacgcgcgatcgcccgcttcgctgccatccggcagcggtctgagcagtagatac
gcggccgtttgcccggtgtgtgggcaattgcggtcccgcagtggcagcggggcccggcgggccgatctggcaatgcctcggcatcgctccgtactctgggcacgagcaacgttcctgtctcgcccggctaaggggcgcgagtctgggagcggacgggt
cggaggtgcgaagtccggcccgttgctctttggtctggtgggaatcctggcaccaatcgggccagaggttccctccgccactcccgacgccccttggggctggtgtgacttggagggccgaagagagccccgccggtgatccggcggggctttgacgtg
cggtcagtgcgtgtgtcggcgagcgatggccacgaggccctggaagccgagcggtccggcgaagtcggcccagtcgcaaccgggctcagcgcagtgggcggaccagccaccgccgttggggtcctggaccaggttcacggtcccctcggtcaggcg
tccgtcgaagtcggtcatggtcggtctcctggtgggtgggggcggggcgccagcacgaagtgccggcgccccgcgggggttggtcgggtcaggcgccgaaccggcgggcggcggcggcgaccaggccgtcggcggcggccatggcgatgatccgct
gcaccgcgggatcggacggatttgcgatggtgtccaactcagtcatggtcgtcctaccggctgctgtgttcagtgacgcgattcctggggtgtgacaccctacgcgacgatggcggatggctgccctgaccggcaatcaccaacgcaaggggaagtcgt
cgctctctggcaaagctccccgctcttccccgtccgggacccgcgcggtcgatccccgcatatatgaagtattcgccttgatccccatcgaattcgtaatcatgtcatagctgtttcctgtgtgaaattgttatccgctcacaattccacacaacatacgag
ccggaagcataaagtgtaaagcctggggtgcctaatgagtgagctaactcacattaattgcgttgcgctcactgcccgctttccagtcgggaaacctgtcgtgccagctgcattaatgaatcggccaacgcgcggggagaggcggtttgcgtattgggcgc
tcttccgcttcctcgctcactgactcgctgcgctcggtcgttcggctgcggcgagcggtatcagctcactcaaaggcggtaatacggttatccacagaatcaggggataacgcaggaaagaacatgtgagcaaaaggccagcaaaaggccaggaacc
gtaaaaaggccgcgttgctggcgtttttccataggctccgcccccctgacgagcatcacaaaaatcgacgctcaagtcagaggtggcgaaacccgacaggactataaagataccaggcgtttccccctggaagctccctcgtgcgctctcctgttccg
accctgccgcttaccggatacctgtccgcctttctcccttcgggaagcgtggcgctttctcatagctcacgctgtaggtatctcagttcggtgtaggtcgttcgctccaagctgggctgtgtgcacgaaccccccgttcagcccgaccgctgcgccttatcc
ggtaactatcgtcttgagtccaacccggtaagacacgacttatcgccactggcagcagccactggtaacaggattagcagagcgaggtatgtaggcggtgctacagagttcttgaagtggtggcctaactacggctacactagaagaacagtatttggta
tctgcgctctgctgaagccagttaccttcggaaaaagagttggtagctcttgatccggcaaacaaaccaccgctggtagcggtggtttttttgtttgcaagcagcagattacgcgcagaaaaaaaggatctcaagaagatcctttgatcttttctacggggtc
tgacgctcagtggaacgaaaactcacgttaagggattttggtcatgagattatcaaaaaggatcttcacctagatccttttggttcatgtgcagctccatcagcaaaaggggatgataagtttatcaccaccgactatttgcaacagtgccgttgatcgtgct
atgatcgactgatgtcatcagcggtggagtgcaatgtcgtgcaatacgaatggcgaaaagccgagctcatcggtcagcttctcaaccttggggttacccccggcggtgtgctgctggtccacagctccttccgtagcgtccggcccctcgaagatgggcc
acttggactgatcgaggccctgcgtgctgcgctgggtccgggagggacgctcgtcatgccctcgtggtcaggtctggacgacgagccgttcgatcctgccacgtcgcccgttacaccggaccttggagttgtctctgacacattctggcgcctgccaaatg
taaagcgcagcgcccatccatttgcctttgcggcagcggggccacaggcagagcagatcatctctgatccattgcccctgccacctcactcgcctgcaagcccggtcgcccgtgtccatgaactcgatgggcaggtacttctcctcggcgtgggacac
gatgccaacacgacgctgcatcttgccgagttgatggcaaaggttccctatggggtgccgagacactgcaccattcttcaggatggcaagttggtacgcgtcgattatctcgagaatgaccactgctgtgagcgctttgccttggcggacaggtggctcaa
ggagaagagccttcagaaggaaggtccagtcggtcatgcctttgctcggttgatccgctcccgcgacattgtggcgacagccctgggtcaactgggccgagatccgttgatcttcctgcatccgccagaggcgggatgcgaagaatgcgatgccgctcg
ccagtcgattggctgagctcatgagcggagaacgagatgacgttggaggggcaaggtcgcgctgattgctggggcaacacgtggagcggatcggggattgtctttcttcagctcgctgatgatatgctgacgctcaatgccgtttggcctccgactaacga
aaatcccgcatttggacggctgatccgattggcacggcggacggcgaatggcggagcagacgctcgtccgggggcaatgagatatgaaaaagcctgaactcaccgcgacgtatcgggccctggccagctagctagagtcgacctgcaggtccccgg
ggatcggtcttgccttgctcgtcggtgatgtacttcaccagctccgcgaagtcgctcttcttgatggagcgcatggggacgtgcttggcaatcacgcgcaccccccggccgttttagcggctaaaaaagtcatggctctgccctcgggcggaccacgccc

atcatgaccttgccaagctcgtcctgcttctcttcgatcttcgccagcagggcgaggatcgtggcatcaccgaaccgcgccgtgcgcgggtcgtcggtgagccagagtttcagcaggccgcccaggcggcccaggtcgccattgatgcgggccagctcg
cggacgtgctcatagtccacgacgcccgtgattttgtagccctggccgacggccagcaggtaggccgacaggctcatgccggccgccgccgccttttcctcaatcgctcttcgttcgtctggaaggcagtacaccttgataggtgggctgcccttcctggt
tggcttggtttcatcagccatccgcttgccctcatctgttacgccggcggtagccggccagcctcgcagagcaggattcccgttgagcaccgccaggtgcgaataagggacagtgaagaaggaacacccgctcgcgggtgggcctacttcacctatcct
gcccggctgacgccgttggatacaccaaggaaagtctacacgaaccctttggcaaaatcctgtatatcgtgcgaaaaaggatggatataccgaaaaaatcgctataatgaccccgaagcagggttatgcagcggaaaagatccgtcgacctgcaggc
atgctaaaccgatacaattaaaggctccttttggagccttttttttggaggcgaactcccttatgcgggtggcgcagaatgccaggacccttgtcattccacgtcaattcatgcgccttttcacctcgtactgtcctgccaaacacaagcaacggaggtacg
catatgtcgaagaagagcaagccgggcgacggtattcgcggcaagggcgtccggGGAcgcttcatgttcatggacgagaccccggaggcggaggccaccacctcgatccgggattccctggccgacctggccgagacgctggaaagcgaagccc
gcaagTAAccagaggcaggtgacggcatgccgtacgagcggatctcccgacgcagactcgcactgtggttgacctggccggccatcaggggcgagttcggcccggcccatcccttcgaggacggtccggatatggcgacggaacccccgtggccc
cggggcgaacctggaatgcgaaatctgcgcacggagttcctcgcctcgCTGcggcgcgccgtcgggaacagcgagggggtcgcggtcgccacctccggcgggctcgactccctcgccgttctggccaacctcatcctcgagcacggtgatcagcg
ccggatcaccgctgtcgccatcgaaatgatcgatgaccgggggaagtccaacatccccgtcgcccggaacctgctcgagggaatcggtctgcggtgcgacctgcggatcgtaccgatctcccggcgaccgacgggggttccggaatggcatgcggaa
gggccacgtctcgacgccctgcccgatgccaaccggcttctggtggaactcgcccacgaggcgggcgcggacgtgctgctcaccggcgacggcgccgacgaactcctgggcagcgggcagtacctgacaccccacCTGtttcgccacggaaac
gtgcgcgcactgccggactactggcgcgatcaccgcggcgagtatcagcccacgaagaaactcgagacgctcgccgtactctcgcccctcctcggccgacgacgccgggcgatgacgtatctggcgtgttcctggccggagctgtgcagcaatcca
ccgcacgacttcctcgccgagccgtaccgaggaatcgccaccgcatggacgaagtcctggatcggttcgctggtacggctccacgaacggcaccatgattcgctggccgtcatggaagcgtggagcgacctctatccacaccaacggctgcgcaca
gccggtctggtgcgcaacgaggaccccttcctggacccggagttcgtcgaacgggcgaagcgactaccgctgcaccggcgctacgacccgagctacccgcacccgtactggcggatgaaatcccaggtgatcaagctgatgcccccgggtgccct
ggcgcatctccccaccgccaaacaaatcttcagccgggccatcgtcagccgtgtggatcccgccgaactgtcggcgaagtatctgctggagagcgggatcctcagcgcccgcgcgctgtcaggagacctggatcccatggtggcgacccgcatcgtc
gaggtcgagcaatggctggccacggcgatggaactcggttacgaactgatcgactaatcacccttgcggagccatggacgaaacctcgaaagtcagccgcgtcaccttccgtactctgctggcctggacgcggggcagtcggcgaaggctgctgttcg
ctctgctctgcgccctggccggatccggactcgggctggcccagcctctgatcgtcaaccggaccatcgaggcggcgacggcggggcggtccgtcgccggactgttgtggctgctcgccgccgtcttcgcaggccaagccgcattgcaggcctggtcg
acgttccggctcgaccaggccggcgagggtgtcgtccagcggctgcgcagggcgatggtcgaccagttgctgtgcttgcgcatggccgtctacgaccgtcaccggctgggcgacctgatctcccgcaccaccaccgacaccaccctgctgcgcgac
gtgctcgccaccgacgtggtgaacgcgatcgtcaactccatcgtcgtggtcggcggcgtgggcatgatgttctggctggatccgatactcgccggggtggtcatggcggtcgtcatcgggtccggagcggtgatggcgctcgtactcaaggggatccgccg
cgagacccaggccgcgcaggaagccaccggctccctcgccgccgagctggagcgcgtcctcagcgccatccgcaccgtgcgcatcaaccgtgccgaggcgcgcgaatccgcggcgctggcccgtagcaccgacgacatctacggtcacaacgt
caaggccgcccgtctgaccgccgtggtctcacccgccatcgaactcgcggtgcagggctcgctgatcgtggtcatcgtcgtcggcctggccagggtcgcgtccggggacatggccgtcggcgagctggtggccttcctcctctacgtgaactacctggc
cgtaccgatggccgggctgttcgacatcgctcccgcactgcaagccgggctcgcggcgctgcagcgcatccaggaggtcctggatctgcccgtcgaaacggaccggctccggacggaccgcccgcagacggaccagttccggaaggagcccgtcg
ccgtcgccgtcgccgcctccgggctcgtgctgcacgacctccggttccgctacaccgatcgcgagattctcaaggggggcgtcgttccgggtgccccgccacagcttcacggccatcgcagggccgtccggggccggcaagagcacgatcttcagcct
gctcggacggttctacgaccccgcggccgggagcatcaccctcgacgggcgcgatgcccacaccgagcacaccacgggcgaatggcggtcccgcatcggcctggtggagcagcattcaccgctgatgtacggcactctgcgcgacaacctcacct
acgcggcgccggacgccgaagaaccgtggctgcaccgggtagtggagatggcgggcttgtccgccttggtgaagcggctgcccgacggcctcgacacccccgtcggggaacacggctcgtcgttgtccggtggcgaacaacagcgcgtggcgatcg
ctcgtgccctgctggcccggccggagatcctgttgctggacgagcccacctcgcacctggacgccgcccatgagcagctcctggcacagaccctgcatctgatctccaaggagtgcacgctgctggtcatcgcccaccggctgtccacgatccggtc
cgccgaccaggtcgtcgtgctggacggcgggcgggtggtcgccgccggcggctacgacgagatacgggaccacctgcccgcgcagcacgcggccgcctgatgaccggtacgggcgacgagcaaccggccgtcctggacagaagaggacccgAt
gaccagtgtggtagccatgacacgcctaacgctccgcccgtaccgcgcgaccgaccggacggccgtcctgacccttgtcaacgccgacaggctcccgggccagccggaagccacacccgacatgctcgcgcaggccCTGcgcggcgagtcgc
ccgtcgacgacaactggtgggccgaactcgacccgccgaccgtacacgtggccaccaccgccgacggcaacggcaacgtcgtgggcgtcgtctcctgcgcactgcgtccgaaagacgacaccgggctgatcctgtgggcccactgccgtgagaa
cgcggccgtggccgaggcgctgatcgcccatgccctggcagagctggcgccccgccccgtcgaagccttccagttcgcgaccgcgctcaccctgggtctggaggcgctgccgatcggccaccggccggtgacccgcgcggcactggaacgcgccg
ggttcaccggcgaacgcctctggcgctacatgcacgccgccctaccgcgccccggcctgccacgggcggacaacatccggatcgtccccgagcccgagcggacgaccgctcggcaactgcaggtgcgcagagacggccggacggtggccgagg
cggtcatcggcgaacccgtccagggcgtcggcgtcctgtggtggatcgaggtcggccccgaagcgcgaaggcagggcctgggacgcgcactactcggctccgccctcgacacgctcaccggtctaggagccactgaagcgattctctacgtcgacg
acgacgaacccccgggggcgaacgcgaccggacggcggccaacgccctgtacgaatccgccggattcaacgaagtcgatcagctgtggatctacgtacggcccaccgtcacgccgtgaagagatcttaaaccgatacaattaaaggctcctttttg
gagccttttttttggaggcgaactcccttatgcgggtggcgcagaatgccaggacccttgtcattccacgtcaattcatgcgccttttcacctcgtactgtcctgccaaacacaagcaacggaggtacggacatgaacaccttcttctcctccgaccaggt
ctcggcgcccgaccgcgtcgcgctctggcacgacgtcatctgccgcagctacgtcccgctgaacatcaccctgaccagcgagcagcccttcatcggcaccgtctccaccggcaacctgggcaccgtccgcatcgccacgtcctcgtccctgcccc
agcagatcacccgcacccgccgcctgatccgccaggacgagcgcgagtacctcatggtcggcgtgcagtccgccggccacgccctcgtgcagcagcacggccgcaccgcccgcgtcggccgcggcggcctggtcttctgggacacccgccacc
cgtacgacatcctcttcccgaccgactggcgcatgagcgtcttccagttcccgcgctactccttcggcttcaccgaggacttcatcggccgcatgaccgccgtcaacgtcggcggcgaccgcggcatcggccgcgtggtctcctccttcatgaccagca
tcaacgacgcgaccgacgccggcgacctggcggaggtcgcctccctccacaacagcgccgtcgacctgctgtccgcggcgatccggaccgagctggccgaccaggccgccgcctccgacggcctgctcgagtgcgtgctcgcctacatccgcca
gaacctggccgacccgaacctgtgcgcctcccagatcgccgccgagcacaacgtctccgtgcggaccctccaccgcctgttctcggccaccggccagggcgtggccgagcacatccgcaacctccgcctcgagcgcatcaagaccgagctggcc
gacccgaccagccggcgctacaccatcagcgccctggcgcgcaagtggggcttcctcgacccgtccaccttctcccgcgcgttcaaggacgcctacggcatcaccgcccgcgagtgggcggcctccgcctccgcctccccgaccgaggtctcgta
gtaaaccgatacaattaaaggctccttttggagccttttttttggagtctagactgcagccgggggtccggcagggccggagcgggcggaaccgtgctctgacctgcggcccgagtttcgtcacgtgacggaatggaaggctgctgcatttcgtcacgtga
cgtatctcggcgagcgactgccgacgccacggcggacacgatcgcctcgcgctggcgccgggcctcgtacgcccgctggcggcaggagcggcggcagtagtcccggctccggccgacgccggattgcttgatctccgagccgcaccaggcgcaga
gcttcgcgccgtcggcgtccctggggtggtggtgctcatggccgacgaccgtacgcggcacgtctcgtagcgaggcgagtcgggcgcgaggtaccgcctgcacgaagtgccggcggggccgacccgggcgagtaatcccaggattactcccgcg
gcttcgaccccggccgccgtcgccgcgtacg
